# Environmental variation is related to morphotype differentiation in a *Brunfelsia* (Solanaceae) complex

**DOI:** 10.64898/2026.04.23.720374

**Authors:** Raquel G. G. Carvalho, Claudio N. Fraga, Mario R. Moura, Leandro L. Giacomin

## Abstract

**Background and aims:** This study combines morphological and environmental data to better understand a *Brunfelsia* (Solanaceae) species complex, aiming to clarify patterns of variation and identify ecological factors that shape morphotype boundaries. Such an approach provides a broader perspective on how organisms respond to environmental gradients and contributes to a more comprehensive understanding of biodiversity.

**Methods:** We analyzed 254 herbarium specimens for 13 morphological traits using univariate and ordination analyses. Climatic and edaphic variables were extracted for all specimens with georeferenced records. To assess habitat suitability and the ecological niche of each predefined morphotype, niche models under present conditions and niche overlap tests were conducted. Ordination analyses were applied to evaluate how environmental predictors explain variation in vegetative and floral traits, whereas supervised multivariate classification was applied to estimate membership probabilities based on morphological and environmental data.

**Key Results:** Integrating morphological, climatic, and edaphic data proved to be a significant approach for refining the species limits within the complex. Although three morphotypes were evaluated, our analyses consistently recovered two well-differentiated groups: the “*capitata*–*hydrangeiformis*” morphotype, allegedly composing a cline, and the ecologically and morphologically distinct “bahia” morphotype. Environmental factors explained more variation in floral than in vegetative traits, with floral traits able to delineate morphotypes more robustly when considered in isolation. The “*capitata*–*hydrangeiformis*” and “bahia” groups also exhibited markedly differences in predicted habitat suitability, with clear separation of the “bahia” morphotype from the others in both geographical and environmental space. Key morphological characters for the taxonomy of *Brunfelsia* partially covaried with environmental variables, particularly floral traits.

**Conclusions:** Our findings support the recognition of “bahia” morphotype as a distinct species to be formally described. This integrative approach contributes to understanding diversification processes in a biodiversity hotspot and reveals previously unrecognized taxonomic diversity within *Brunfelsia*, a genus that harbors many rare and narrow-endemic taxa.

## INTRODUCTION

Species are increasingly understood as testable evolutionary hypotheses, rather than fixed entities, whose boundaries can be corroborated or refuted as new evidence accumulates (Pante *et al*. 2015). Under the unified concept, species correspond to independently evolving metapopulation lineages (De Queiroz 2007), implying that species delimitation must be grounded in explicit hypotheses about lineage divergence. This perspective is particularly relevant for species complexes, in which extensive overlap in morphological attributes obscures clear taxonomic boundaries and challenges traditional typological approaches (de Souza *et al*. 2023). In such complexes, speciation is best investigated through the joint evaluation of multiple axes of variation, including morphology, geography, and ecology, across the full geographic range of the taxon (Wood and Nakazato 2009; Cadena and Cuervo 2010).

Morphological characters remain central to taxonomic practice, particularly when subtle but consistent phenotypic differences accumulate among closely related lineages (Zapata 2014). However, morphology alone may be insufficient when phenotypic variation appears to be continuous or environmentally structured, making geographic and ecological information essential for identifying lineage limits and diversification processes (Cadena and Cuervo 2010). In plants, this challenge is especially pronounced, as continuous clines in vegetative and reproductive traits often track environmental gradients, potentially obscuring species boundaries when morphology is considered in isolation (Wood and Nakazato 2009; Alcántara-Ayala *et al*. 2020; He *et al*. 2022). Such patterns are common in widespread plant taxa distributed across heterogeneous landscapes, such as in the Atlantic Forest biodiversity hotspot (Zachos and Habel 2011). The environmental complexity of this region provides an ideal setting for testing whether morphological differentiation reflects adaptive divergence, phenotypic plasticity, or the presence of previously unrecognized evolutionary lineages.

Within this context, *Brunfelsia* L. (Solanaceae: Petunieae) emerges as a compelling model system for testing hypotheses of species limits. This medium-sized shrub genus comprises 49 species distributed across two centers of diversity in South America and the Antilles (Plowman 1998; Filipowicz and Renner 2012). While South American species can also occur in the Cerrado and Amazon, much of the genus’ diversity is concentrated in the Atlantic Forest, including 14 species, 11 of which are endemic to Brazil (Ribeiro-Silva and Knapp 2020).Most species of *Brunfelsia* have narrow, medium-length corolla tubes, a nectar guide at the tube opening, exhibit diurnal anthesis with floral color change, and are pollinated by butterflies (Plowman 1998; Filipowicz and Renner 2012). Floral traits in the genus are known to be closely linked to pollination syndromes, suggesting a degree of ecological specialization that may promote reproductive isolation and lineage divergence (Filipowicz and Renner 2012). Despite this ecological relevance, overlapping morphological variation among populations has historically motivated the recognition of infraspecific taxa within several species of the genus. This raises questions about whether such variation reflects environmentally driven divergence along ecological gradients, the occurrence of a single highly variable lineage (e.g., an ochlospecies), or the presence of distinct evolutionary lineages occupying partially overlapping environmental niches and/or distribution (Cronk 1998; Braby *et al*. 2012; Patten 2015). When such lineages are overlooked because they exhibit few or no diagnostic morphological differences, they may be considered ‘cryptic’ species (e.g. Renner 2020). Consideration of these alternative hypotheses suggest that *Brunfelsia hydrangeiformis* (Pohl) Benth. may represent a species complex.

Against this broader backdrop, *Brunfelsia hydrangeiformis* represents a particularly illustrative case. This locally rare shrub is distributed along eastern Brazil, from São Paulo to southern Bahia states, where it occurs mainly in well-preserved forest formations associated with water bodies. It is readily recognized in the field by its large leaves and conspicuous, many-flowered inflorescences of purple flowers that turn white during anthesis. Two subspecies are currently recognized, – *B. hydrangeiformis* subsp. *hydrangeiformis* and *B. hydrangeiformis* subsp. *capitata* (Benth.) Plowman, – distinguished primarily by variation in leaf morphology, pedicel length, and calyx characters (Plowman 1998). Since the last comprehensive taxonomic treatment of the genus (Plowman 1998), the substantial increase in herbarium material has revealed a broader and more structured pattern of morphological variation within the *B. hydrangeiformis* complex. Notably, specimens from the northernmost portion of the distribution exhibit a striking combination of traits contrasting with southern populations. These geographic and morphological differences, coupled with the species’ broad environmental distribution, suggest that current taxonomic boundaries may mask environmentally structured evolutionary lineages occupying distinct regions of environmental space.

In this study, we use an integrative framework that combines morphometric analyses, geographic distribution, and ecological niche characterization to assess the taxonomic circumscription of the *Brunfelsia hydrangeiformis* species complex. Specifically, we assess whether patterns of morphological variation correspond to environmentally structured lineages with distinct ecological niches, and whether this evidence supports the maintenance of the currently recognized subspecies or the recognition of other taxa. By explicitly linking morphological differentiation to ecological space, this study aims to refine species boundaries within *Brunfelsia* and to contribute to a broader understanding of how environmental heterogeneity shapes diversification in tropical plant lineages, particularly within the Atlantic Forest hotspot.

## MATERIALS AND METHODS

### Taxon sampling

Two hundred and fifty-four (254) herbarium specimens were analyzed (see Data Availability) through databased images from SpeciesLink (https://specieslink.net), GBIF (https://www.gbif.org), Reflora (https://reflora.jbrj.gov.br) and JABOT (https://jabot.jbrj.gov.br). For the CEPEC (acronyms following Thiers, continuously updated) collection, where no images were available in the databases, photographs were taken directly during a visit to the herbarium. In addition, floral cards were made from analyzed exsiccates in the most relevant collections for the species (CEPEC, BHCB, and RB), to allow measuring floral attributes more precisely, considering the species very congested inflorescence. For the organization of the specimens to be analyzed, classification into morphotypes was carried out, following Plowman’s taxonomic treatment for the species *B. hydrangeiformis* (Plowman 1998). Therefore, specimens corresponding to the subspecies presented in Plowman were classified as morphotypes “*capitata*” and “*hydrangeiformis*”, and the morphotype “bahia” was assigned to specimens witha corolla tube twice the length of the calyx, distinguishing them from the other morphotypes (Fig. 1). All data acquired, as well as all scripts of the analysis described below are publicly available in a Zenodo repository (Carvalho *et al*. 2026).

**Fig. 1.**
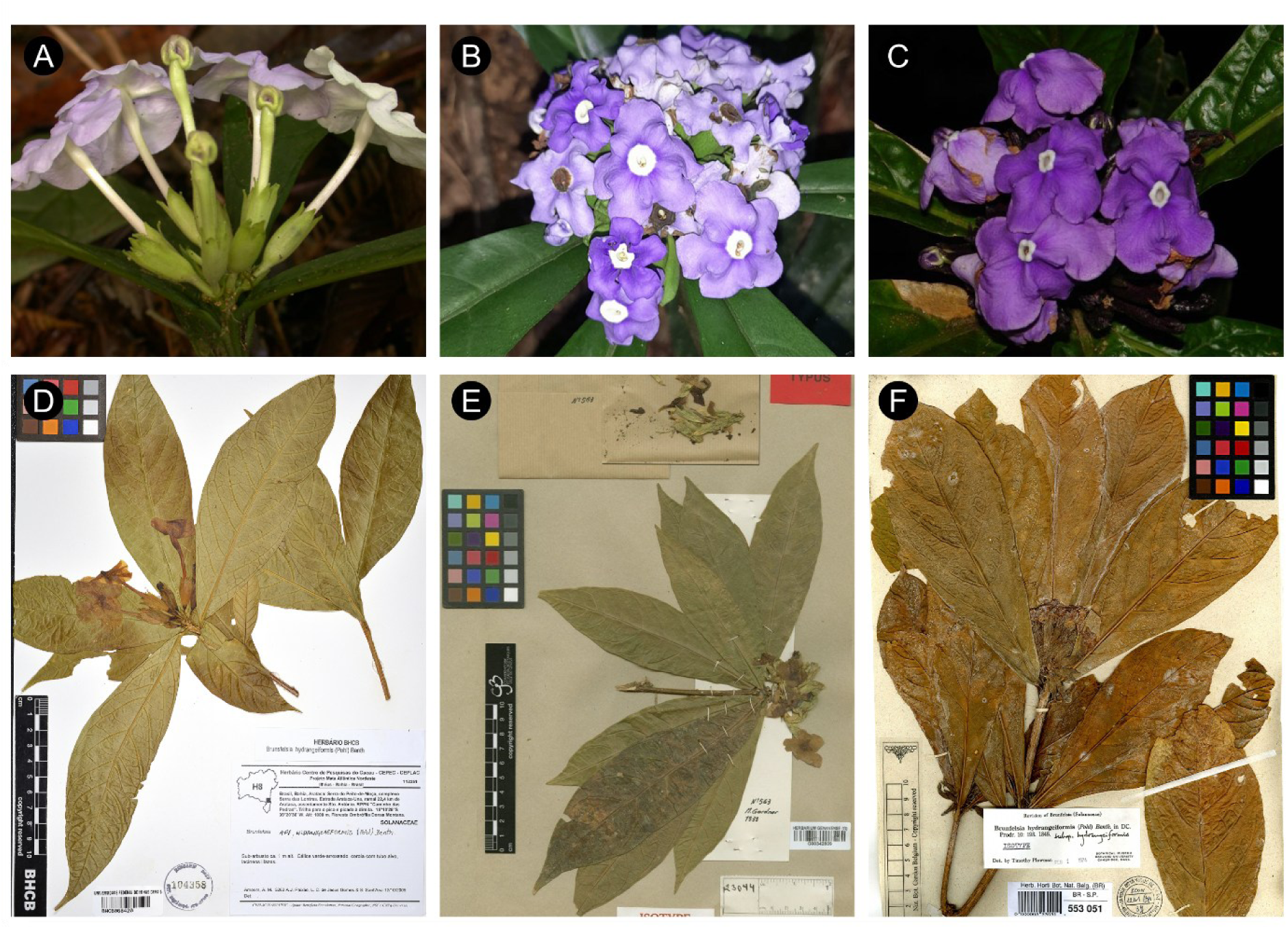
*Brunfelsia hydrangeiformis* morphotypes. (A-C) Flowering plant images taken in their habitat for each morphotype: (A) “bahia” morphotype with the considerably longer corolla tube; (B) “*capitata”* morphotype and (C) “*hydrangeiformis*” morphotype. Photos provided by A.M.Amorim, J.M.A.Braga and C.N.Fraga, respectively. (D-F) Dried specimens for each morphotype: (D) BHCB104358 A.M.Amorim, 5263; (E) G00342809 M.Gardner, 563; (F) BR0000005530519 J.B.E.Pohl, s.n.; in the (E) picture is possible to note that narrower leaves of “*capitata*” morphotype when compared to “*hydrangeiformis”*, a character used to separate them in Plowman (1998).

### Morphometric analysis

Images of the exsiccates and floral cards (all standardized with scale bars), were analyzed and all measurements were obtained using ImageJv.1.54g (https://imagej.net/ij/; Schneider *et al*. 2012). Thirteen (13) morphological characters (Fig. 2) were measured, including six vegetative and seven reproductive traits. For each specimen, at least three measurements were taken for each character. To visualize the main morphometric differences and variation between morphotypes, violin plots with values of statistical metrics were created from the morphometric dataset. To assess statistical significance, ANOVA was applied to normally distributed data, while the non-parametric Kruskal-Wallis test was used for data that did not meet normality assumptions (Supplementary Data Fig. S1). Morphometric traits were examined with a multivariate approach, including Principal Component Analysis (PCA), to summarize trait variation, and Canonical Variate Analysis (CVA), in order to identify putative groups, implemented in the R v. 4.5.0 software (R Core Team 2025).

**Fig. 2.**
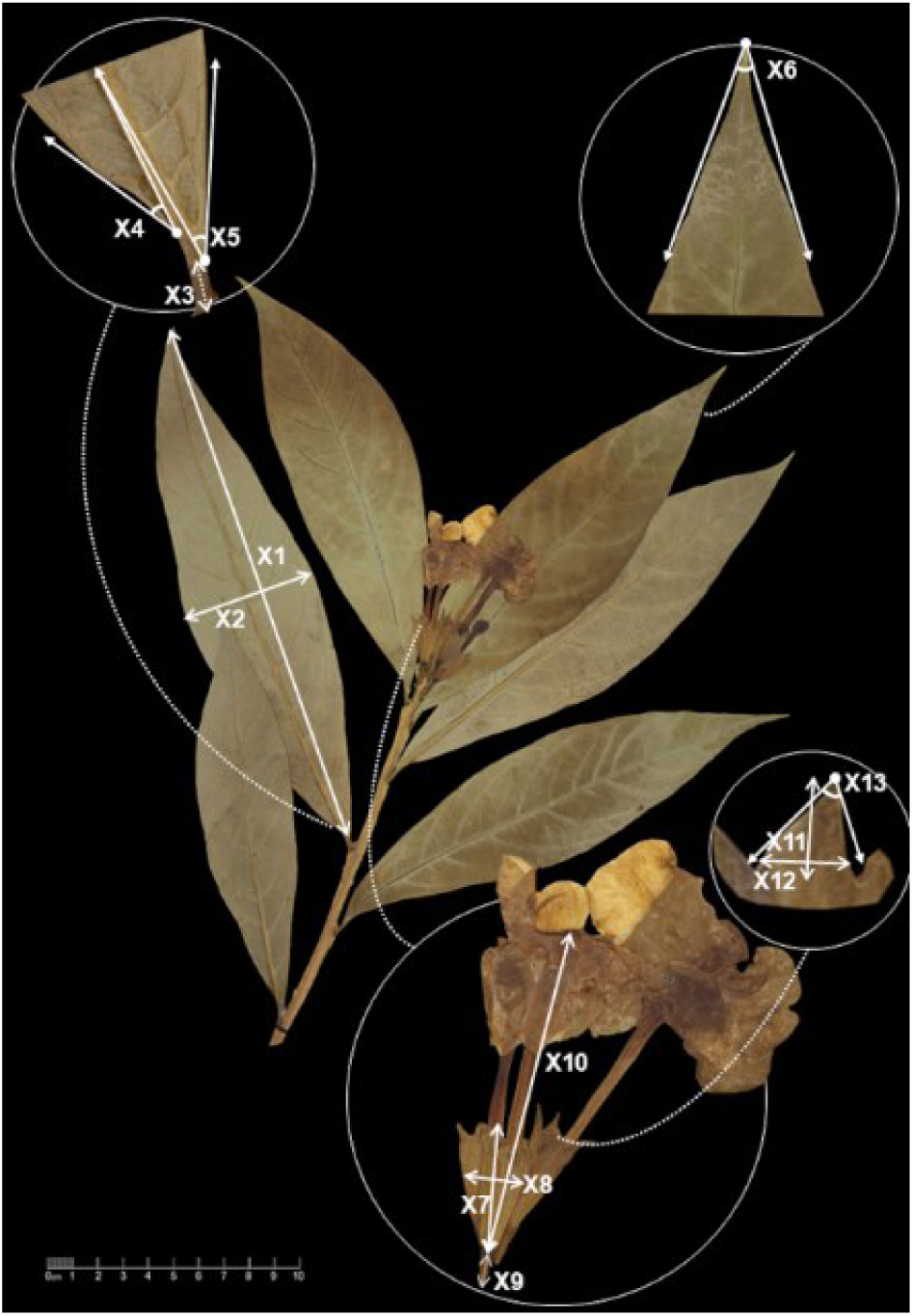
Vegetative and reproductive floral characters used in morphometric analysis. X1 = Lamina length (cm)/Lam_len; X2 = Lamina width (at widest) (cm)/Lam_wid; X3 = Petiole length (cm)/Pet_len; X4 = Lamina base angle (right side) (°)/Lam_bas_rang; X5 = Lamina base angle (left side) (°)/Lam_bas_lang; X6 = Lamina apex angle (°)/Lam_ape_ang; X7 = Calyx length (cm)/Cal_len; X8 = Calyx width (at widest) (cm)/Cal_wid; X9 = Pedicel length (cm)/Ped_len; X10 = Corolla tube length (cm)/Cor_tub_len; X11 = Calyx teeth length (cm)/Cal_tee_len; X12 = Calyx teeth width (at widest) (cm)/Cal_tee_wid; X13 = Calyx teeth angle (°)/Cal_tee_ang.

### Ecological niche modelling

Geographical coordinates of each herbarium specimens were obtained from voucher metadata. All occurrence records were cleaned using the *CoordinateCleaner* package (Zizka *et al*. 2019), removing invalid coordinates, duplicates, marine records, and conversion errors. For each morphotype, the calibration area was defined using the buffered minimum convex polygon (BMCP) method with a 1000 km buffer. Within each calibration area, pseudo-absences were generated at twice the number of presences to avoid strongly imbalanced datasets, following the environmentally constrained method to ensure ecological plausibility (Engler *et al*. 2004). Spatial thinning was then applied to ensure only one occurrence per raster cell was kept (Kramer-Schadt *et al*. 2013), retaining a total of 92 occurrence records.

To represent environmental space, we used the 19 bioclimatic variables from CHELSA v2.1 (Karger *et al*. 2017) and SoilGrids v2.0, depth 5–15 cm (Poggio *et al*. 2021) databases downloaded at 30 arc-sec of spatial resolution and degraded to 2.5 arc min (∼5km). To reduce multicollinearity among predictors, we computed the variance inflation factor (VIF) for the full set of environmental variables, iteratively removing the variable with the highest VIF until all remaining variables had VIF values ≤ 5 (Kutner *et al*. 2005). Overall, 14 variables were retained for the subsequent procedure (Supplementary Data Table S2).

Since the choice of algorithm can affect ENM predictions, we computed an ensemble of projections (Araújo and New 2007) based on five methods: Generalized Linear Models (using linear terms only); Generalized Additive Models (using smooth terms with three dimensions); Gradient Boosting Model (automatically tuned using the following parameter variation of learning rate [0.1, 0.5, 1], number of trees [50, 100, 250, 500], and number of leaves in the terminal nodes [1, 3, 5, 7, 9]); Neural Networks (hyperparameters tuned based on combinations of size of hidden layers [2, 4, 6, 8, 10] and weights for parameter decay [0.001, 0.05, 0.1, 0.5, 1, 3]); and Random Forests (with the entry parameter automatically tuned [values 1, 3, or 5] through growing 1000 trees; (Breiner *et al*. 2015)).

To validate each ENM, we applied a 10-fold cross-validation, with 90% of the data used for model calibration and 10% reserved for testing in each iteration, ensuring independence between model calibration and evaluation. Model performance was evaluated using the Sorensen similarity index, calculated between observed and predicted presences, and indicated due to its independence of species prevalence (Leroy *et al*. 2018). The ensemble model for each species was computed as the average weighted habitat suitability across algorithms, with weights given by the Sørensen index calculated for each algorithm. All ENM operations were performed in R using the *flexsdm* R package (Velazco *et al*. 2022). Raw data and code are available through Data Availability.

### Niche Overlap Analysis

We assessed niche overlap, using equivalency and similarity tests, directly in the environmental space defined by the current climatic (CHELSA v2.1) and edaphic (SoilGrids 2.0, depth 5–15 cm) (Poggio *et al*. 2021) conditions of the selected environmental variables (Supplementary Data Table S2). To reduce the dimensionality and avoid multicollinearity, we applied a Principal Component Analysis (PCA) to the climatic and edaphic predictors separately, retaining the first two axes to perform niche overlap analyses. We measured niche equivalency and niche similarity between all pairs of morphotypes using the Schoener’s D statistic (Schoener 1968; Warren *et al*. 2008), which ranges from 0 (complete niche divergence) to 1 (complete niche identity), computed separately for climatic and edaphic dimensions.

The niche equivalency test evaluates whether two species occupy identical positions within niche space, by pooling occurrences and repeatedly randomizing them into datasets matching the original sample sizes to generate a null distribution of niche overlap, against which the observed overlap is compared, independently of the environmental background (Warren *et al*. 2010). The niche similarity test assesses whether niche overlap exceeds random expectations by randomly shifting the occurrence points of one species within environmental space and recomputing Schoener’s D across multiple iterations to generate a null distribution (Warren *et al*. 2010). Because estimates of niche similarity depend on the environmental conditions considered available to each taxon, the background environmental space was defined as the union of the ENM calibration areas (i.e., 1000 km buffered minimum convex polygon) across the three morphotypes. We used 1000 iterations to build the null distribution of D values across climatic and edaphic spaces, allowing evaluation of whether morphotype niches are statistically equivalent or more similar than expected by chance. Computations were performed in the *ecospat* R package (Di Cola *et al*. 2017).

### Environmental correlates of morphotype variation

To identify the environmental drivers underlying ecological and morphological differentiation among morphotypes, we quantified the magnitude and direction of predictor effects on habitat suitability models and evaluated associations between environmental gradients and morphometric traits.

For ecological niche models, predictor importance was estimated using a permutation-based approach, in which each predictor was randomly permuted 100 times while all remaining predictors were held constant. Importance was quantified as the reduction in model performance (drop in Sørensen similarity index) following permutation, with larger decreases indicating greater predictor importance (Velazco *et al*. 2022). Effect direction was characterized from partial dependence plots (PDPs) generated for each predictor, morphotype, and modelling algorithm. Pearson’s correlation coefficient was calculated between the predictor gradient and the corresponding model-predicted habitat suitability. Positive values indicate that suitability generally increases along the predictor gradient, whereas negative values indicate the opposite. Correlations close to zero indicate no overall monotonic trend and may reflect complex non-linear responses, particularly in machine-learning algorithms.

To assess whether environmental gradients were associated with morphological variation, pairwise Pearson correlation tests were performed between morphometric traits and environmental predictors. Unlike the ecological niche models, which were fitted separately for each morphotype, these analyses were designed to detect continuous trait–environment relationships across the entire species complex, rather than differences among predefined groups. P-values were adjusted for multiple comparisons using the Benjamini–Hochberg false discovery rate (FDR) correction (Benjamini & Hochberg 1995). All analyses were conducted in the R environment. Operations were performed using the *flexsdm* (Velazco *et al*. 2022) and *stats* package in R (R Core Team 2025).

### Discriminant Analysis of Principal Components

To assess specimen classification and estimate membership probabilities among morphotypes, we performed a Discriminant Analysis of Principal Components (DAPC) using predefined morphotype assignments. The analysis integrated morphological measurements with the selected climatic and edaphic predictors extracted for each georeferenced specimen. Because DAPC requires complete datasets, missing values were first imputed using an iterative PCA-based procedure implemented in the *missMDA* R package (Josse and Husson 2016). The optimal number of principal components used for imputation was estimated by leave-one-out cross-validation, allowing missing entries to be reconstructed from the multivariate covariance structure of the dataset. A Principal Component Analysis (PCA) was then performed on the completed dataset, and the resulting PC scores were used as input for the DAPC. The number of retained PC axes limited to the maximum allowed given the sample size and number of groups, and the number of discriminant functions set to the theoretical maximum (number of groups – 1). Posterior membership probabilities were estimated for each specimen and used to assign individuals to the morphotype with the highest probability. Operations were performed using the *adegenet* R package (Jombart *et al*. 2010).

## RESULTS

Across all analyses, the three *Brunfelsia hydrangeiformis* morphotypes showed consistent patterns of differentiation in both morphology and environmental space, although the magnitude of differentiation varied among analytical approaches. Floral characters contributed most strongly to morphological discrimination, whereas vegetative traits showed extensive overlap. Ecological niche models and niche overlap analyses further revealed contrasting environmental associations among morphotypes, while environmental variables explained a modest but detectable proportion of morphological variation, particularly in floral traits.

### Morphological differentiation among morphotypes

Morphological differentiation among the three morphotypes was driven primarily by floral rather than vegetative characters. Canonical variate analysis (CVA) based on vegetative traits showed a high degree of overlap among the morphotypes, indicating limited discriminatory power despite the first canonical axis explaining 53.2% of variation (Fig. 3A). However, floral traits produced a much clearer separation, with the first canonical axis explaining 73.8% of variation and distinguishing the three morphotypes, with the “bahia” group showing highly reduced overlap (Fig. 3B).

**Fig. 3.**
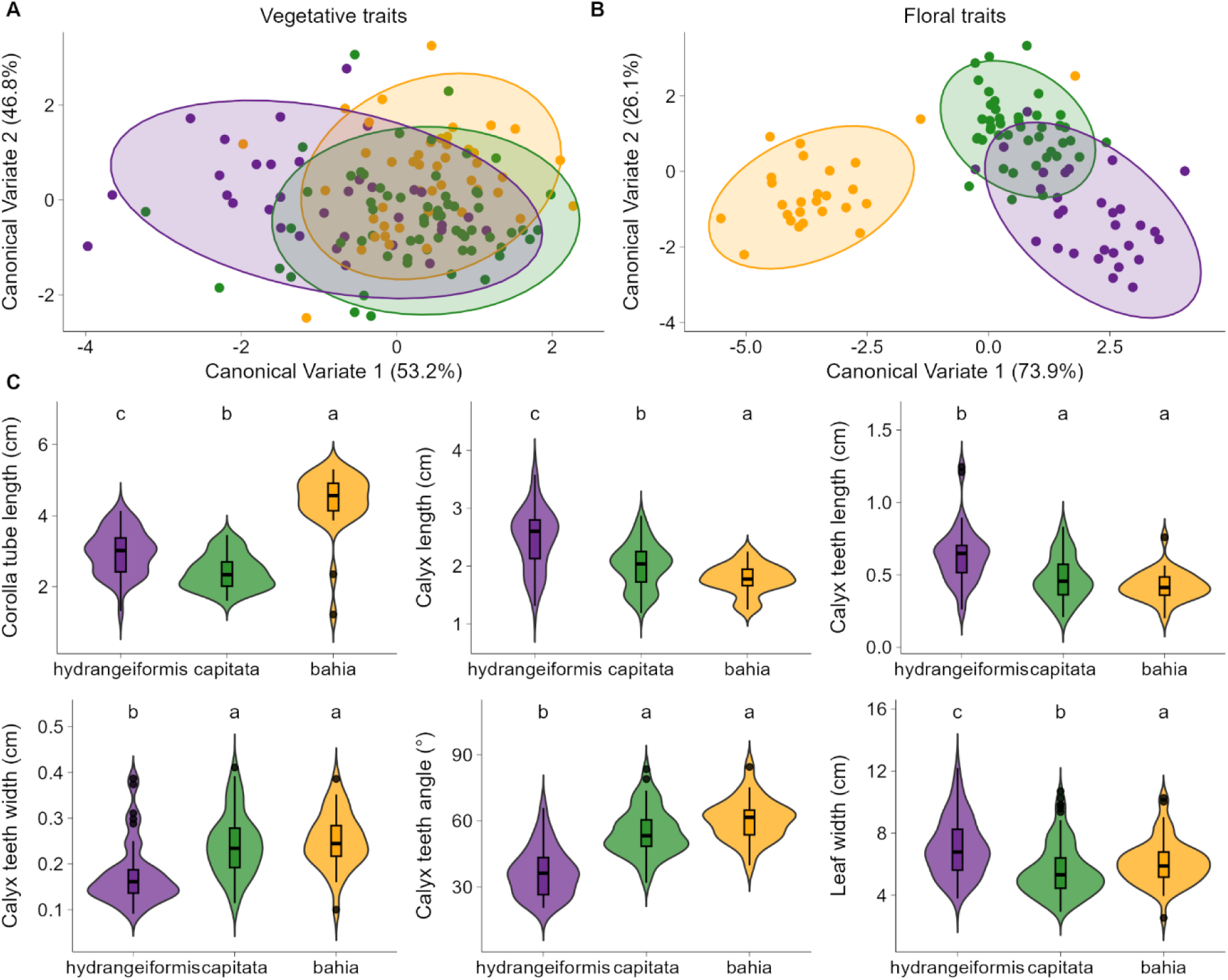
Floral traits provide stronger morphological discrimination among *Brunfelsia hydrangeiformis* morphotypes than vegetative traits. Canonical Variate Analysis (CVA) based on (A) vegetative and (B) floral traits. Points represent individual specimens and ellipses indicate the 95% confidence interval for each morphotype. (C) Distributions of the six morphological traits contributing most strongly to morphotype differentiation. Violin plots represent kernel density estimates, and boxplots indicate the median and interquartile range. Different letters (a,b,c) denote significant differences among morphotypes based on ANOVA or Kruskal–Wallis tests followed by post hoc comparisons (P < 0.05). See Supplementary Data Fig. S1 for univariate results for the complete set of morphometric traits.

Univariate analyses corroborated these multivariate patterns. The “bahia” morphotype differing from at least one other morphotype across all floral traits (ANOVA or Kruskal–Wallis tests, all P ≤ 0.002; Fig. 3C; Supplementary Data Fig. S1). The “bahia” morphotype presented the longest corolla tubes and pedicels. Calyx length increased progressively from “bahia” to “*capitata*” and reached its highest values in “*hydrangeiformis*”. Vegetative traits showed comparatively weaker differentiation. Although leaf width differed among all morphotypes (P ≤ 0.015, Fig. 3C), the distributions of other vegetative traits broadly overlapped between the “bahia” and the other morphotypes. However, “*hydrangeiformis*” and “*capitata*” also differed in petiole length and leaf apex angle (Supplementary Data Fig. S1).

### Ecological niche models and geographic patterns

The ecological niche models showed high predictive performance for the three morphotypes, with ensemble models showing Sorensen similarity values of 1.00 for “bahia”, 0.949 for “*capitata*”, and 0.940 for “*hydrangeiformis*”. Predicted habitat suitability revealed distinct geographic patterns across the Brazilian Atlantic Forest (Fig. 4). The “bahia*”* morphotype was associated with highly suitable habitats in the northern Atlantic Forest, along the eastern Brazilian coast, a region predicted to be largely unsuitable for the other two morphotypes. In contrast, “*hydrangeiformis*” and “*capitata*” morphotypes showed broadly overlapping distributions concentrated in the southeastern Atlantic Forest, with suitable habitat for the “*capitata*” morphotype extending farther inland, whereas “*hydrangeiformis”* remained more closely associated with coastal regions.

**Fig. 4.**
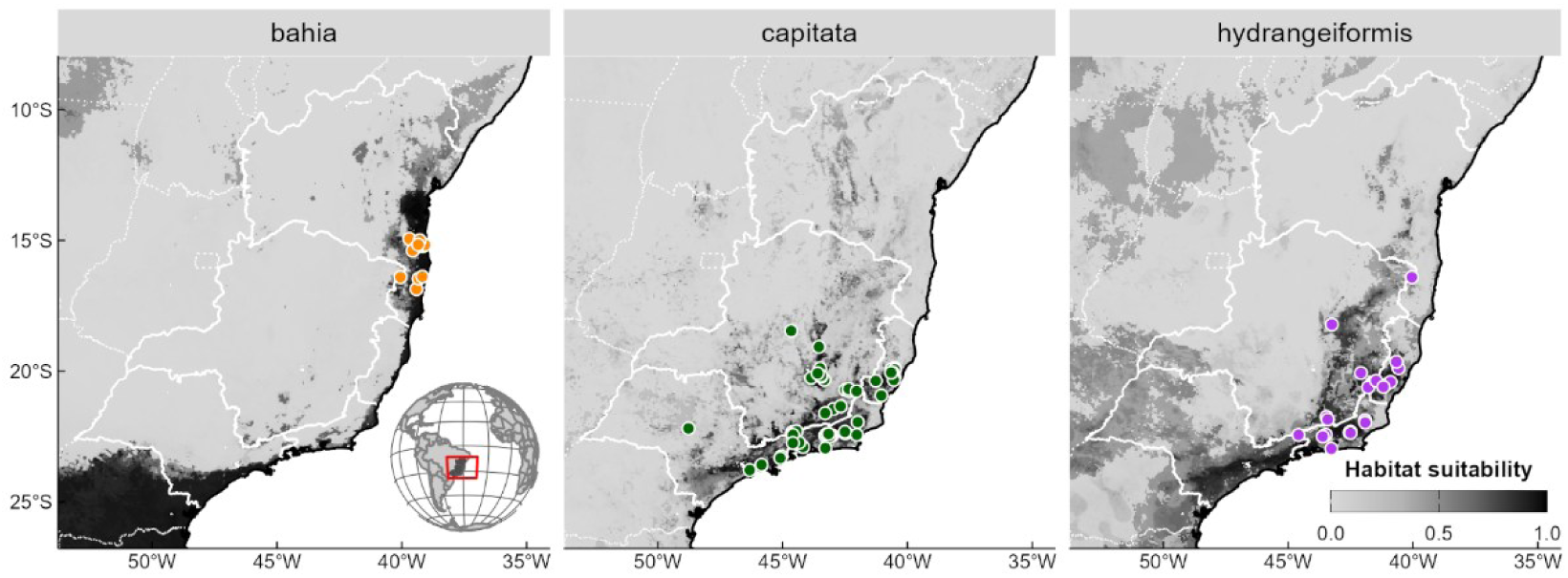
Predicted habitat suitability for three morphotypes recognized in *Brunfelsia hydrangeiformis* (Pohl) Benth. Ensemble ecological niche models under current conditions for the morphotypes “bahia”, “*capitata*”, and “*hydrangeiformis*” across the Brazilian Atlantic Forest. Colored circles represent occurrence records used for model calibration. Habitat suitability ranges from low (light grey) to high (black), with darker shades indicating increasingly suitable environmental conditions. Solid contour lines highlight the states where specimens were recorded: Bahia, Espírito Santo, Minas Gerais, Rio de Janeiro and São Paulo.

### Climatic and edaphic niche differentiation

Pairwise niche overlap analyses revealed contrasting patterns of climatic and edaphic differentiation among the three morphotypes (Fig. 5A). In both environmental spaces, the “bahia” morphotype exhibited lower niche equivalency than the expected by chance when compared with either “*capitata*” or “*hydrangeiformis*” morphotypes (all P < 0.001), indicating that its occupied niche is environmentally distinct. Between “*capitata*” and “*hydrangeiformis*”, we found niche equivalency within the expected by chance in both climatic (P = 0.133) and edaphic (P = 0.375) spaces. Regarding niche similarity, no pair of morphotypes exhibited niches more similar than expected by chance in the climatic space, but all pairs showed higher than expected similarity in the edaphic space.

**Fig. 5.**
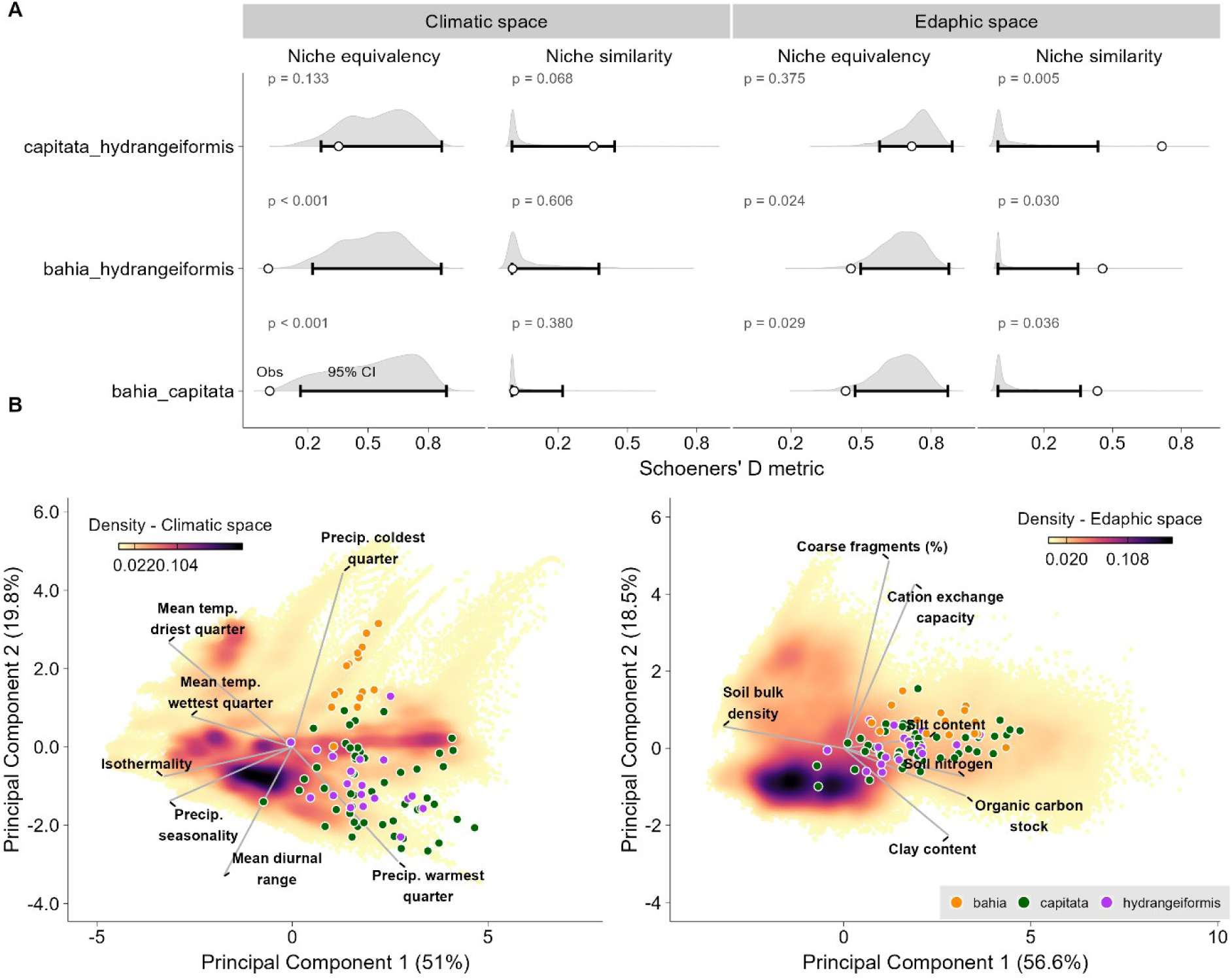
Climatic and edaphic niche differentiation among morphotypes of the *Brunfelsia hydrangeiformis* complex. (A) Pairwise niche equivalency and niche similarity tests in climatic and edaphic space based on Schoener’s D. Grey violins represent the null distributions generated by randomization, white circles indicate the observed overlap, horizontal bars represent the central null distribution, and P values correspond to one-tailed randomization tests. (B) Principal component analysis (PCA) of climatic space. Background shading represents the density of environmental conditions available across the study region, colored points correspond to occurrence records of each morphotype, and vectors indicate the loadings of climatic variables.

### Environmental correlates of morphotype variation

Environmental predictors showed contrasting contributions to ecological niche models among morphotypes (Fig. 6A, Supplementary Data Table S3). Although the relative importance of individual predictors varied among modelling algorithms, “*capitata*” and “*hydrangeiformis*” displayed broadly similar predictor rankings and response directions, whereas “bahia” exhibited a more distinct environmental profile. Across algorithms, cation exchange capacity was consistently the most important predictor for “bahia” and showed positive relationship with habitat suitability, whereas its overall effect was negative in “*capitata*” and “*hydrangeiformis”*. For the morphotypes “*capitata”*, the organic carbon stock was most influential, showing an overall positive effect on habitat suitability, whereas for “*hydrangeiformis*”, the mean temperature of the driest quarter emerged as the most important predictor, affecting habitat suitability negatively. Despite these differences, isothermality and mean diurnal range consistently exhibited a negative association with habitat suitability across all three morphotypes.

**Fig. 6.**
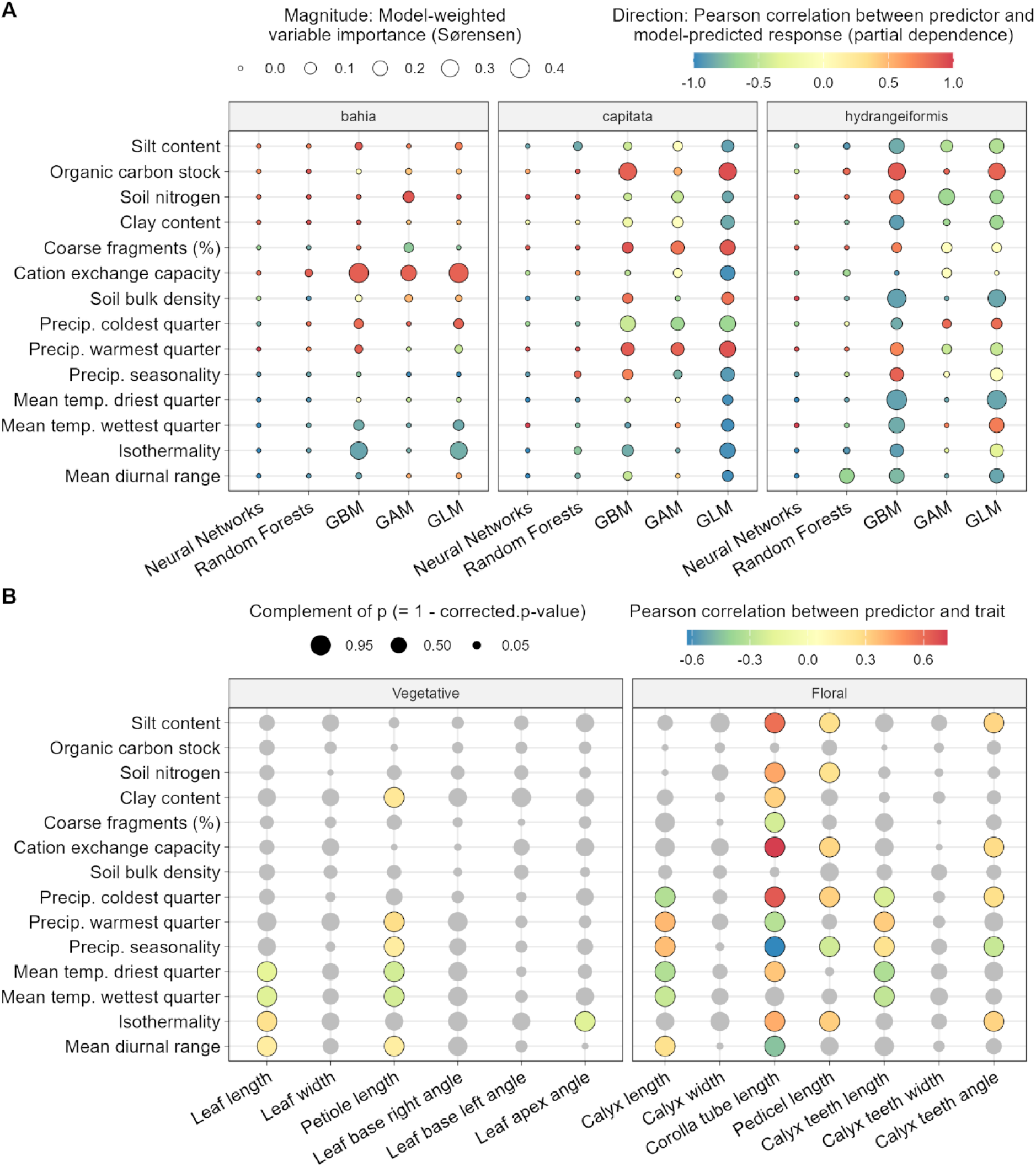
Environmental drivers of ecological niche variation and morphological trait clines in *Brunfelsia hydrangeiformis* complex. (A) Bubble chart summarizing the relative importance and overall direction of environmental predictor effects in ecological niche models for the three morphotypes. Bubble size represents model-weighted predictor importance, while color indicates the overall direction of relationship between predictor values and habitat suitability. (B) Bubble chart summarizing trait–environment associations across the *Brunfelsia hydrangeiformis* complex. Bubble size reflects the complement of the corrected p-value (i.e., statistical significance), and color denotes the strength and direction of Pearson correlations coefficients between environmental predictors and vegetative and floral traits. Gray circles represent non-significant correlations after Benjamini–Hochberg correction. See Supplementary Data Tables S3-S4 for complete numerical results.

Trait–environment correlations further supported this pattern (Fig. 6B). Vegetative traits exhibited relatively few significant associations and generally weak effect sizes (Supplementary Data Table S4), with leaf length showing its strongest positive association with isothermality (Spearman *r* = 0.26) and petiole length with precipitation during the warmest quarter (*r* = 0.25). In contrast, floral traits showed more numerous and stronger relationships with environmental predictors. Among edaphic variables, cation exchange capacity and silt content showed consistent positive associations with multiple floral traits, particularly corolla tube length, pedicel length, and calyx teeth angle. The strongest positive correlation was observed between corolla tube length and precipitation during the warmest quarter (r = 0.38), whereas mean temperature of the driest quarter showed consistent negative associations with calyx length, calyx teeth length, and corolla tube length (all *r* ≈ −0.32 to −0.33). Overall, these results indicate that floral morphology exhibit stronger and more coordinated responses to environmental gradients than vegetative morphology.

### Integrated classification of morphotypes

The discriminant analysis combining morphological, climatic, and edaphic information revealed a strong separation among morphotypes (Fig. 7). The DAPC retained ten principal components (80.2% of the total variance) and two discriminant functions, correctly reclassifying 87.8% of the specimens. Classification accuracy was highest for “bahia” (98.3%), followed by “*capitata*” (91.5%), whereas “*hydrangeiformis*” showed substantially lower assignment success (71.2%). Posterior membership probabilities derived from the DAPC showed that specimens assigned to “bahia” formed a discrete and well-supported group, with virtually all individuals classified with near-complete membership probabilities. In contrast, “*capitata*” and “*hydrangeiformis*” exhibited a continuum of assignment probabilities. Although most specimens were assigned to their a priori morphotype, posterior probabilities gradually declined near the boundary between the two groups, with increasing probabilities of alternative assignment.

**Fig. 7.**
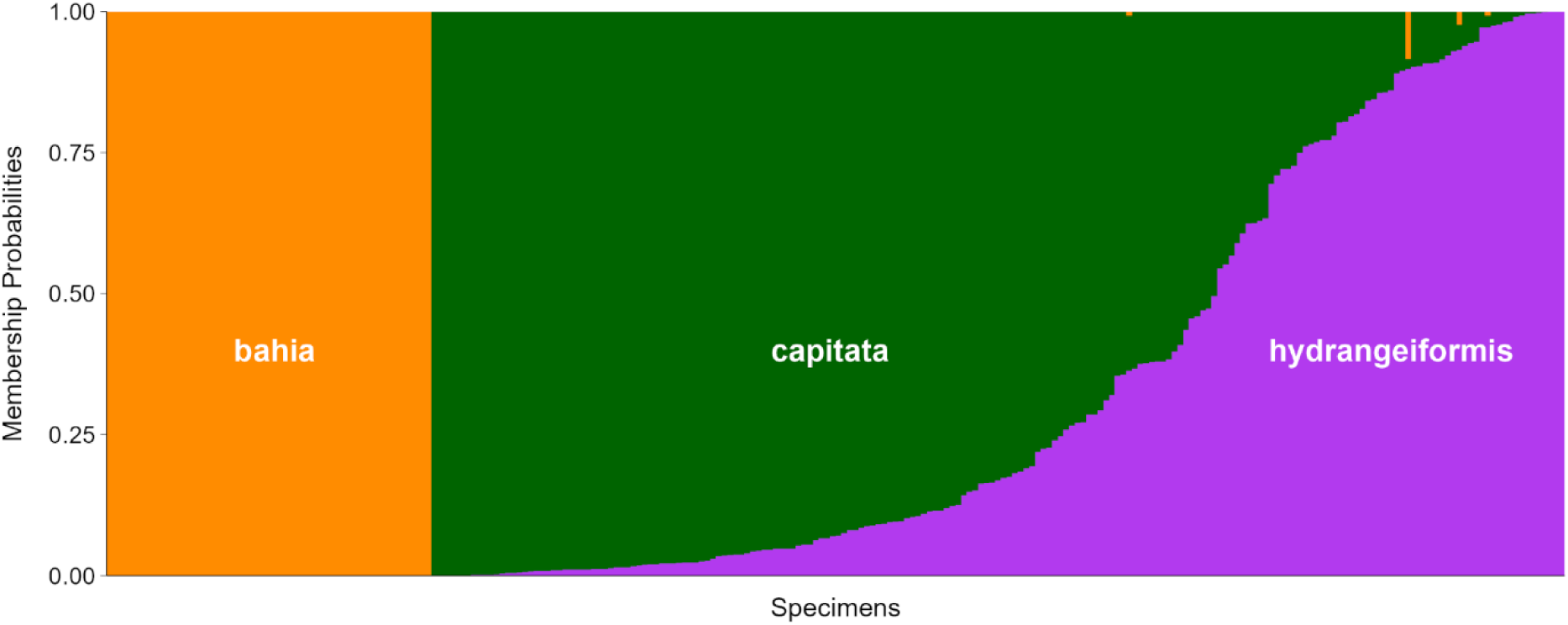
Integrated classification of *Brunfelsia hydrangeiformis* morphotypes based on morphology and environmental data. Posterior membership probabilities estimated by Discriminant Analysis of Principal Components (DAPC) for each specimen using combined morphometric, climatic, and edaphic predictors. Each vertical bar represents one individual, order by its assigned morphotype and posterior membership probability. Specimens with bars dominated by a single color indicate high classification confidence, whereas multicolored bars identify individuals with intermediate membership probabilities and greater overlap among morphotypes. See Supplementary Data Table S5 for details on individual membership probabilities.

## DISCUSSION

The integration of morphological, ecological niche, and environmental analyses consistently recovered two major groups within *Brunfelsia hydrangeiformis* complex comprises: a well-defined “bahia” group, and a second group comprising the partially overlapping morphotypes “*capitata*” and “*hydrangeiformis*”, previously recognized as subspecies by Plowman (1998). Although morphological data alone enabled the identification of key diagnostic traits among these groups, the incorporation of environmental variables substantially improved the resolution and robustness of their delimitation. It is notable that the greater availability of specimens for analysis since Plowman (1998) was key to establishment of a priori hypothesis to be tested and allowed the proposition of the morphotypes tested here. Plowman (1998) had a single specimen from Bahia at hand, which was seemingly not enough to allow the recognition of substantial variation. This reinforces the importance of fieldwork and digitization of data and images for biodiversity comprehension (Soltis 2017; Nelson and Ellis 2018; Swain and Chakraborty 2024).

### Variation in morphologic and environmental space

Individual morphometric analyses play a crucial role in structuring the morphological diversity and variation within plant species and serve as a valuable tool for the identification and delimitation of taxonomic entities (Buitrago Aristizábal *et al*. 2020; Teixeira *et al*. 2022; da Silva-Oliveira *et al*. 2025). In some studies, taxonomic delimitation was successfully achieved using morphometric analyses alone (e.g. Martínez-Flores *et al*. 2025; da Silva-Oliveira *et al*. 2025), as the initial ordination analyses accounted for over half of the total data variation. In contrast, this level of explanatory power was not observed in the present study, although variation was captured when traits were analyzed separately, with the most significant differences between “bahia” and the other morphotypes associated with floral traits (Fig. 3B).

When examining the environmental findings (Fig. 5) alongside niche modeling results (Fig. 4), it becomes evident that the “bahia” morphotype is ecologically distinct, suggesting adaptive pressures and unique factors influencing the occurrence of this metapopulation. Such segregation aligns with the framework of ecological speciation in plants, where divergent selection across heterogeneous habitats fosters reproductive isolation and lineage diversification (Givnish 2010). Cosacov *et al*. (2013), through an analysis of morphological trait variation in populations of *Calceolaria polyrhiza* Cav. (Calceolariaceae), demonstrated that corolla size exhibits a geographic pattern congruent with vegetative traits, being primarily shaped by climatic gradients, as well as by pollinator interactions. This evidence is consistent with the meta–analysis of Caruso *et al*. (2019), which highlights that abiotic factors can impose substantial selective pressures on floral traits. In this context, the “bahia” morphotype may represent an incipient stage of ecological speciation, with its distribution shaped by localized environmental factors that reduce gene flow with other populations and promote adaptive divergence.

Studies integrating ecological and molecular datasets (Reck-Kortmann *et al*. 2017; Backes *et al*. 2023; Chase *et al*. 2024) provide fundamental insights into the delimitation of species complex. Augsten *et al*. (2025) exemplified this integrative framework by combining ecological niche modeling with plastid DNA markers to investigate a complex within *Nicotiana* sect. *Alatae* Goodsp. (Solanaceae). Extending such molecular approaches to the *Brunfelsia hydrangeiformis* complex, in conjunction with morphological and ecological evidence, would markedly improve the resolution of morphotype boundaries and enable rigorous testing of phylogeographic barriers—as demonstrated by Soares *et al*. (2025) in their population genomic assessment of species delimitation within *Petunia* (Solanaceae). Moreover, the incorporation of molecular data reinforces the application of De Queiroz’s (2007) unified species concept, thereby yielding more robust and comprehensive evidence for species delimitation, picturing in what stage of the speciation continuum the species might be.

### Putative influence of the environment on morphological variation

The integration of morphological and ecological datasets was pivotal for delineating the taxonomic circumscription in this study. This integrative approach is widely adopted for such purposes (Ceolin and Miotto 2012; Zapata 2014; García *et al*. 2020; Teixeira *et al*. 2022; Augsten *et al*. 2025), as it enables the construction of a robust analytical framework that enhances statistical reliability in taxon delimitation. Teixeira *et al*. (2022) demonstrated that variation in floral shape and size between the two analyzed morphotypes was significantly correlated with climatic variables, suggesting that these differences are likely driven by processes of species diversification and speciation, leading to the formal recognition of a new species.

Correlation analysis between morphological traits and environmental predictors demonstrated that floral characteristics yielded the most significant associations (Fig. 6B), thereby establishing them as diagnostic features across morphotypes, both when examined independently (Fig. 3) and within integrated datasets (Fig. 6B). Notably, the “bahia” morphotype—characterized by individuals with more elongate corolla tubes (see Fig. 3C)—exhibited a completely distinct ecological space (Fig. 5) and was influenced by key environmental predictors that diverged from those shaping the other morphotype models (Fig. 6A). Consequently, this morphotype can be regarded as the most differentiated among those analyzed, where variation seems to be structured. In *Brunfelsia*, corolla traits—including tube length, coloration, and scent—are pivotal for understanding evolutionary patterns and shifts in pollination strategies (Filipowicz and Renner 2012). Species within the section *Franciscea* Pohl (sensu Plowman 1998), which encompasses *B. hydrangeiformis*, typically exhibit shorter and narrower corolla tubes, with a characteristic constriction at mouth, central nectar guides, and color changes during anthesis, features that collectively characterize a psychophilous pollination syndrome (Vogel 1954; Cocucci 1995; Plowman 1998). Although this pollination mechanism has been described in the genus (Cocucci 1995) and a dichotomy between psychophilous and sphingophilous syndromes was used to recognize putative lineages (see Vogel 1954; Plowman 1998) floral biology studies for the genus are scarce. Since variation in corolla tube length may reflect adaptive responses associated with shifts in pollinator specialization (Summers *et al*. 2015), it is an interesting topic that deserves further investigation in the genus. The putative association of a pollination shift and a speciation event make the “bahia*”* morphotype an ideal model for testing such a hypothesis, since it presents a much longer corolla tube compared to the other morphotypes associated to *B. hydrangeiformis*. Investigating variation in scent among these morphotypes, as well as other species of section *Franciscea* also deserves attention (see Cocucci 1995), since it can be related to a compartmentalization of pollinators among the populations.

The only study on *Brunfelsia* available in the literature that investigated the relationship between corolla traits and pollinators within a phylogenetic context was that of Filipowicz *et al*. (2012), based on morphometric data obtained from literature. Research on diverse plant groups has revealed associations between variation in corolla tube length and shifts in pollinator assemblages (Cohen 2012; Newman and Johnson 2021; McCarren *et al*. 2023). Newman and Johnson (2021) demonstrated that differences in floral tube length across two populations of *Erica junonia* Bolus (Ericaceae) correspond to variation in proboscis length among nemestrinid fly species. Through pollinator exclusion experiments, they showed that the short-tubed variety can produce seeds in the absence of pollinators, whereas the long-tubed variety depends fully on pollinators. This case thus exemplifies a change in corolla tube length directly linked to a shift in pollinators — an investigation that would be particularly compelling to conduct in populations of these *Brunfelsia* morphotypes.

### Structuring of groups within morphotypes

CVA results indicated that the morphotype groups were more clearly structured when analyzing only floral data, but there was still a high degree of overlap when analyzing vegetative traits, with no significant separation between them and no clear positioning within a quadrant (see Fig. 3A-3B). Meanwhile, discriminant analysis revealed partial overlap between “*capitata”* and “*hydrangeiformis*” morphotypes, while “bahia” morphotype showed high membership probability (Fig. 7). These results demonstrate that while the discriminant analysis captures a strong axis of divergence, the probability-based assignments showed finer-scale patterns of uncertainty and admixture, providing a more nuanced view of morphological structuring among the groups of morphotypes. In some studies, the discriminant analysis used to infer membership probabilities is shown to be highly recommended, revealing an accurate method to this purpose (Buitrago Aristizábal *et al*. 2020). Since the results indicate a high degree of morphological and ecological overlap between “*hydrangeiformis*” and “*capitata*” morphotypes, these two subspecies will be synonymized and treated as a single taxon in a future taxonomic revision. The morphological characters that once were used to separate them might indicate an initial stage of separation within the speciation continuum, what should be tested in the future with appropriate methods. Given the taxonomic path forward, investigating gene flow among these populations would help elucidating the degree of separation between them and what stage of the speciation continuum (Stankowski and Ravinet 2021) would they really be.

The ecological niche models and overlap test results support the perspective that while “*hydrangeiformis*” and “*capitata*” are ecologically closer, “bahia*”* is differentiated by its unique environmental associations, with a restricted niche, consistent with its narrower distribution and potential ecological specialization. The distinct ecological niche identified for the “bahia*”* morphotype may be closely related to the unique biogeographic and historical context of southern Bahia state of Brazil. This region, historically referred to as the “Hileia Baiana” (Faria *et al*. 2021), corresponds to the so-called “Bahia refuge” (Carnaval and Moritz 2008; Martins 2011) which represent portions of the Atlantic Forest that remained climatically stable throughout the Pleistocene and Holocene. Such long-term stability has been associated with reduced extinction rates and the accumulation of species over time (Monteiro Lopes *et al*. 2016). Furthermore, the geomorphological heterogeneity of the region, including Quaternary coastal plains, Oligocene “tabuleiros,” and inland mountain ranges (Barbosa *et al*. 2003), together with high levels of forest conservation and extensive remnants (Ribeiro *et al*. 2009) contribute to its exceptional floristic diversity (Faria *et al*. 2021), which is still being described (e.g. Goldenberg *et al*. 2020; Torres *et al*. 2020; Morales *et al*. 2021; Gaem *et al*. 2024). These historical and environmental factors likely underlie the ecological distinctiveness of the “bahia*”* morphotype, reinforcing the role of southern Bahia as both a hotspot of woody plant diversity and a center of endemism within the Atlantic Forest in Brazil, deserving further conservation initiatives.

## CONCLUSION

The integrated evidence presented here demonstrates that the *Brunfelsia hydrangeiformis* complex is composed of two well-differentiated groups: the “*capitata–hydrangeiformis*” morphotypes and the ecologically distinct “bahia*”* morphotype, that can be distinguished by its considerably longer corolla tube. While morphological traits alone provided partial resolution for the recognition of the “bahia” morphotype, the incorporation of ecological niche modeling and environmental predictors substantially enhanced the robustness of its delimitation. The “bahia*”* morphotype exhibits a restricted distribution and unique ecological associations that are consistent with its occurrence in the historically stable and floristically rich landscapes of southern Bahia, representing a distinct species to be formally described. Meanwhile “*capitata*-*hydrangeiformis*” morphotypes may represent a single taxonomic entity, rather than separate subspecies, and will be formally synonymized in a future taxonomic revision. Key morphological features seem to covary with environmental variables, pointing to a putative ecological speciation that is worth testing. Equally important is the need for studies in floral biology of *Brunfelsia*, particularly those addressing the relationship between corolla tube length and shifts in pollinator assemblages in southern Bahia state, in Brazil. These findings highlight the importance of combining morphometric and ecological approaches to uncover hidden diversity, refine species boundaries, and better understand the evolutionary and ecological processes shaping plant lineages in biodiversity hotspots such as the Atlantic Forest.

## SUPPLEMENTARY DATA

Supplementary data are available at Annals of Botany online and consist of the following. Table S1: Summary of descriptive quantitative characters measured on 254 specimens of Brunfelsia *hydrangeiformis* divided into three morphotypes: *hydrangeiformis*; *capitata* and *bahia*. Table S2: Variation Inflation Factor (VIF) for predictor variables retained for ecological niche modeling. Table S3: Predictor importance and direction of environmental effects for ecological niche models. Table S4: Pearson correlations between morphometric traits and environmental predictors across the *Brunfelsia hydrangeiformis* complex. Table S5: Membership probabilities of each individual to its assigned morphotype according to DAPC analysis. Figure S1: Violin plots for all morphometric characters analyzed.

## FUNDING

This work is funded by Coordenação de Aperfeiçoamento de Pessoal de Nível Superior – Brazil (CAPES) (88887.948805/2024-00), through the master’s scholarship provided to the first author, linked to the postgraduate program at the Botanical Garden of Rio de Janeiro (JBRJ). LLG and CNF are funded by CNPq (grants 422191/2021-3, 408914/2023-8, and 404996/2024-8). In addition, CNF is supported by FAPERJ (grant no. 201280/2022) and CNPq-PPBio (grant no. 441250/2023-8), MRM is supported by CNPq (grant no 409197/2024-6; part of the National Institute of Science and Technology (INCT) in Ecology, Evolution, and Biodiversity Conservation).

## ACKNOWLEDGEMENTS

We are grateful for the support of the curators of the aforementioned herbaria. We also thank the researchers A.M. Amorim, and J.M.A. Braga for the field images provided, as well as researchers C. Turchetto, M. Trovó and E. Pessoa for their suggestions and considerations throughout the course of the research. The authors acknowledge the funding provided by CAPES, CPNq and FAPERJ, and all the support from the National School of Tropical Botany (ENBT) - JBRJ throughout the master’s project.

## DATA AVAILABILITY

The scripts and data used in this project are publicly available in a Zenodo repository (Carvalho *et al*. 2026; doi: 10.5281/zenodo.21863887).

## Notes

https://zenodo.org/records/21863887

### Competing Interest Statement

The authors have declared no competing interest.

### Summary of Updates

This version of the manuscript has been revised to update the methods, figures, discussion, and supplementary material, incorporating suggestions made by reviewers to date. Significant changes were made to the niche modeling analysis; the RDA and PCA analyses were removed while the CVA analysis was retained; the structure of the Materials and Methods section was reorganized; the discussion was expanded; and the figures were rearranged.

