## Supplementary Data for "Environmental variation is related to morphotype differentiation in a *Brunfelsia* (Solanaceae) complex"

**This PDF file includes**

Supplementary Tables S1-S5

Supplementary Figure S1

**TABLE S1.** Summary of descriptive quantitative characters measured on 254 specimens of *Brunfelsia hydrangeiformis* divided into three morphotypes: *hydrangeiformis*; *capitata* and *bahia*. (5 percentile−) mean ± standard deviation (−95 percentile). CH= character code.

| CH | Character | *hydrangeiformis* | *capitata* | *bahia* |
|---|---|---|---|---|
| Lam_len | Lamina length (cm) | (13,678−) 21,419 ± 4,969 (−30,01) | (13,429−) 19,076 ± 4,466 (−28,100) | (14,829−) 20,412 ± 3,909 (−25,499) |
| Lam_wid | Lamina width (at widest) (cm) | (4,54−) 7,168 ± 1,786 (−10,521) | (3,465−) 5,586 ± 1,64 (−8,741) | (4,150−) 6,135 ± 1,426 (−8,567) |
| Pet_len | Petiole length (cm) | (0,327−) 0,648 ± 0,203 (−0,996) | (0,315−) 0,545 ± 0,164 (−0,884) | (0,376−) 0,563 ± 0,143 (−0,811) |
| Lam_ape_ang | Lamina apex angle (º) | (47,863−) 69,835 ± 14,178 (−93,019) | (43,568−) 61,680 ± 13,193 (−85,772) | (44,893−) 58,874 ± 11,097 (−77,073) |
| Lam_bas_rang | Lamina base angle (right side) (º) | (23,961−) 35,938 ± 11,524 (−50,677) | (22,099−) 35,371 ± 8,871 (−48,610) | (19,840−) 34,848 ± 9,843 (−49,555) |
| Lam_bas_lan | Lamina base angle (left side) (º) | (25,262−) 38,227 ± 10,251 (−54,617) | (22,793−) 41,021 ± 11,861 (−62,026) | (23,316−) 37,138 ± 9,694 (−55,155) |
| Cal_len | Calyx length (cm) | (1,857−) 2,610 ± 0,410 (−3,219) | (1,388−) 1,994 ± 0,404 (−2,604) | (1,250−) 1,774 ± 0,280 (−2,235) |
| Cal_wid | Calyx width (at widest) (cm) | (0,435−) 0,618 ± 0,176 (−0,942) | (0,406−) 0,689 ± 0,170 (−1,018) | (0,477−) 0,713 ± 0,188 (−1,014) |
| Cal_tee_len | Calyx teeth length (cm) | (0,353−) 0,625 ± 0,200 (−0,896) | (0,237−) 0,471 ± 0,149 (−0,709) | (0,297−) 0,430 ± 0,100 (−0,562) |
| Cal_tee_wid | Calyx teeth width (at widest) (cm) | (0,106−) 0,171 ± 0,061 (−0,280) | (0,14−) 0,616 ± 3,492 (−0,365) | (0,151−) 0,252 ± 0,063 (−0,346) |
| Cal_tee_ang | Calyx teeth angle (º) | (23,220−) 35,422 ± 10,660 (−55,879) | (37,447−) 54,474 ± 10,896 (−73,167) | (42,448−) 59,664 ± 9,985 (−74,343) |
| Cor_tub_len | Corolla tube length (cm) | (2,259−) 3,108 ± 0,572 (−3,967) | (1,829−) 2,415 ± 0,482 (−3,260) | (3,334−) 4,465 ± 0,710 (−5,269) |
| Ped_len | Pedicel length (cm) | (0,312−) 0,530 ± 0,189 (−0,872) | (0,296−) 0,481 ± 0,130 (−0,709) | (0,359−) 0,620 ± 0,159 (−0,861) |

**TABLE S2. Variation Inflation Factor (VIF) for predictor variables retained for ecological niche modeling.** Climatic data were extracted from CHELSA, whereas edaphic data were obtained from SoilGrids.

| Variable acronym | Variable description | Unit | VIF |
|---|---|---|---|
| bio02 | Mean Diurnal Range (Mean of monthly max temperature − min temperature) | °C | 2.6 |
| bio03 | Isothermality (BIO2 / BIO7 × 100) | % | 2.23 |
| bio08 | Mean Temperature of Wettest Quarter | °C | 4.8 |
| bio09 | Mean Temperature of Driest Quarter | °C | 4.83 |
| bio15 | Precipitation Seasonality (Coefficient of Variation) | % | 1.93 |
| bio18 | Precipitation of Warmest Quarter | mm | 1.89 |
| bio19 | Precipitation of Coldest Quarter | mm | 1.96 |
| bdod | Bulk Density of Fine Earth | kg $dm^{-3}$ | 2.93 |
| cec | Cation Exchange Capacity | cmol(c) $kg^{-1}$ | 2.58 |
| cfvo | Volumetric Fraction of Coarse Fragments (>2 mm) | $cm^3$ $dm^{-3}$ (vol%) | 2.52 |
| clay | Clay Content | g $kg^{-1}$ | 1.56 |
| nitrogen | Total Nitrogen | g $kg^{-1}$ | 3.4 |
| ocs | Organic Carbon Stock | t $ha^{-1}$ | 4.41 |
| silt | Silt Content | g $kg^{-1}$ | 1.37 |

**TABLE S3. Predictor importance and direction of environmental effects for ecological niche models.** Model-weighted variable importance was quantified as the reduction in predictive performance (drop in Sørensen similarity) following permutation of each predictor. Predictor direction was estimated as the Pearson correlation between predictor values and model-predicted habitat suitability derived from partial dependence plots. Positive values indicate an overall increase in predicted suitability with increasing predictor values, whereas negative values indicate the opposite. Correlations close to zero indicate the absence of an overall monotonic response and may reflect complex non-linear relationships.

| | Weighted Importance | | | Spearman R | | |
|---|---|---|---|---|---|---|
| **Algorithm/Predictor** | **bahia** | **capitata** | **hydrangeiformis** | **bahia** | **capitata** | **hydrangeiformis** |
| **GAM** | | | | | | |
| Cation exchange capacity | 0.237 | 0.038 | 0.052 | 0.857 | 0.000 | 0.000 |
| Clay content | 0.000 | 0.085 | 0.010 | 0.443 | 0.000 | -0.635 |
| Coarse fragments (%) | 0.049 | 0.153 | 0.066 | -0.716 | 0.778 | 0.000 |
| Isothermality | 0.000 | 0.000 | 0.000 | -0.826 | -0.849 | -0.340 |
| Mean diurnal range | 0.000 | 0.000 | 0.000 | 0.547 | 0.394 | -0.866 |
| Mean temp. driest quarter | 0.000 | 0.000 | 0.000 | -0.488 | 0.000 | -0.866 |
| Mean temp. wettest quarter | 0.000 | 0.000 | 0.000 | -0.853 | 0.635 | 0.740 |
| Organic carbon stock | 0.006 | 0.023 | 0.003 | 0.446 | 0.520 | 0.849 |
| Precip. coldest quarter | 0.000 | 0.146 | 0.038 | 0.853 | -0.635 | 0.827 |
| Precip. seasonality | 0.000 | 0.030 | 0.003 | -0.984 | -0.809 | 0.000 |
| Precip. warmest quarter | 0.000 | 0.139 | 0.052 | -0.472 | 0.860 | -0.471 |
| Silt content | 0.000 | 0.050 | 0.109 | 0.736 | 0.000 | -0.563 |
| Soil bulk density | 0.019 | 0.001 | 0.000 | 0.505 | -0.636 | -0.860 |
| Soil nitrogen | 0.084 | 0.097 | 0.253 | 0.905 | -0.470 | -0.636 |
| **GBM** | | | | | | |
| Cation exchange capacity | 0.423 | 0.002 | 0.000 | 0.871 | -0.516 | -0.930 |
| Clay content | 0.000 | 0.049 | 0.180 | 0.893 | -0.190 | -0.902 |
| Coarse fragments (%) | 0.000 | 0.080 | 0.055 | 0.804 | 0.917 | 0.731 |
| Isothermality | 0.310 | 0.093 | 0.146 | -0.868 | -0.836 | -0.899 |
| Mean diurnal range | 0.004 | 0.034 | 0.200 | -0.871 | -0.479 | -0.786 |
| Mean temp. driest quarter | 0.000 | 0.001 | 0.463 | 0.000 | -0.466 | -0.883 |

| Algorithm/Predictor | Weighted Importance | | | Spearman R | | |
|---|---|---|---|---|---|---|
| | bahia | capitata | hydrangeiformis | bahia | capitata | hydrangeiformis |
| Mean temp. wettest quarter | 0.067 | 0.002 | 0.244 | -0.839 | -0.892 | -0.839 |
| Organic carbon stock | 0.001 | 0.330 | 0.322 | 0.000 | 0.862 | 0.889 |
| Precip. coldest quarter | 0.048 | 0.237 | 0.088 | 0.817 | -0.459 | -0.837 |
| Precip. seasonality | 0.000 | 0.067 | 0.154 | -0.735 | 0.718 | 0.860 |
| Precip. warmest quarter | 0.024 | 0.147 | 0.138 | 0.869 | 0.887 | 0.705 |
| Silt content | 0.016 | 0.027 | 0.217 | 0.902 | -0.448 | -0.839 |
| Soil bulk density | 0.010 | 0.064 | 0.375 | 0.000 | 0.792 | -0.883 |
| Soil nitrogen | 0.000 | 0.021 | 0.177 | 0.859 | -0.452 | 0.778 |
| **GLM** | | | | | | |
| Cation exchange capacity | 0.420 | 0.199 | 0.000 | 0.859 | -0.961 | 0.000 |
| Clay content | 0.000 | 0.161 | 0.164 | 0.464 | -0.850 | -0.635 |
| Coarse fragments (%) | 0.000 | 0.219 | 0.052 | -0.730 | 0.896 | 0.000 |
| Isothermality | 0.301 | 0.228 | 0.148 | -0.827 | -0.958 | -0.340 |
| Mean diurnal range | 0.002 | 0.080 | 0.184 | 0.549 | -0.968 | -0.866 |
| Mean temp. driest quarter | 0.000 | 0.064 | 0.406 | -0.500 | -0.964 | -0.866 |
| Mean temp. wettest quarter | 0.070 | 0.120 | 0.203 | -0.853 | -0.971 | 0.740 |
| Organic carbon stock | 0.001 | 0.330 | 0.291 | 0.465 | 0.935 | 0.849 |
| Precip. coldest quarter | 0.053 | 0.246 | 0.074 | 0.856 | -0.640 | 0.813 |
| Precip. seasonality | 0.000 | 0.166 | 0.133 | -0.995 | -0.915 | 0.000 |
| Precip. warmest quarter | 0.022 | 0.254 | 0.125 | -0.467 | 0.917 | -0.471 |
| Silt content | 0.014 | 0.113 | 0.198 | 0.737 | -0.887 | -0.563 |
| Soil bulk density | 0.007 | 0.113 | 0.324 | 0.504 | 0.791 | -0.860 |
| Soil nitrogen | 0.000 | 0.087 | 0.161 | 0.926 | -0.857 | -0.636 |
| **Neural Networks** | **0.000** | **0.000** | **0.000** | **-0.145** | **-0.192** | **-0.170** |
| Cation exchange capacity | 0.000 | 0.000 | 0.000 | 0.807 | -0.553 | -0.751 |
| Clay content | 0.000 | 0.000 | 0.000 | 0.831 | -0.118 | -0.517 |
| Coarse fragments (%) | 0.000 | 0.000 | 0.000 | -0.667 | 0.902 | 0.870 |
| Isothermality | 0.000 | 0.000 | 0.000 | -1.000 | -1.000 | -1.000 |

| Algorithm/Predictor | Weighted Importance | | | Spearman R | | |
|---|---|---|---|---|---|---|
| | bahia | capitata | hydrangeiformis | bahia | capitata | hydrangeiformis |
| Mean diurnal range | 0.000 | 0.000 | 0.000 | -1.000 | -1.000 | -1.000 |
| Mean temp. driest quarter | 0.000 | 0.000 | 0.000 | -0.981 | -0.998 | -1.000 |
| Mean temp. wettest quarter | 0.000 | 0.000 | 0.000 | -0.993 | 0.998 | 1.000 |
| Organic carbon stock | 0.000 | 0.000 | 0.000 | 0.682 | 0.570 | -0.435 |
| Precip. coldest quarter | 0.000 | 0.000 | 0.000 | -0.824 | -0.651 | -0.648 |
| Precip. seasonality | 0.000 | 0.000 | 0.000 | -0.914 | -0.903 | -0.881 |
| Precip. warmest quarter | 0.000 | 0.000 | 0.000 | 0.996 | 0.935 | 0.937 |
| Silt content | 0.000 | 0.000 | 0.000 | 0.767 | -0.865 | -0.816 |
| Soil bulk density | 0.000 | 0.000 | 0.000 | -0.558 | -0.960 | 0.981 |
| Soil nitrogen | 0.000 | 0.000 | 0.000 | 0.816 | 0.958 | 0.886 |
| **Random Forest** | 0.020 | 0.065 | 0.234 | -0.025 | -0.037 | -0.255 |
| Cation exchange capacity | 0.020 | 0.000 | 0.010 | 0.871 | 0.609 | -0.626 |
| Clay content | 0.000 | 0.000 | 0.000 | 0.934 | 0.137 | -0.769 |
| Coarse fragments (%) | 0.000 | 0.000 | 0.000 | -0.829 | 0.911 | 0.860 |
| Isothermality | 0.000 | 0.017 | 0.006 | -0.884 | -0.730 | -0.925 |
| Mean diurnal range | 0.000 | 0.001 | 0.204 | -0.922 | -0.823 | -0.662 |
| Mean temp. driest quarter | 0.000 | 0.000 | 0.000 | -0.932 | -0.920 | -0.880 |
| Mean temp. wettest quarter | 0.000 | 0.000 | 0.000 | -0.854 | -0.772 | -0.678 |
| Organic carbon stock | 0.000 | 0.000 | 0.009 | 0.914 | 0.895 | 0.826 |
| Precip. coldest quarter | 0.000 | 0.000 | 0.000 | 0.798 | -0.787 | -0.116 |
| Precip. seasonality | 0.000 | 0.010 | 0.000 | -0.860 | 0.845 | -0.474 |
| Precip. warmest quarter | 0.000 | 0.000 | 0.000 | 0.697 | 0.923 | 0.814 |
| Silt content | 0.000 | 0.036 | 0.006 | 0.732 | -0.848 | -0.922 |
| Soil bulk density | 0.000 | 0.000 | 0.000 | -0.941 | -0.811 | -0.827 |
| Soil nitrogen | 0.000 | 0.000 | 0.000 | 0.921 | 0.850 | 0.816 |

**TABLE S4. Pearson correlations between morphometric traits and environmental predictors across the *Brunfelsia hydrangeiformis* complex.** Correlation coefficients (r) quantify the strength and direction of the association between each morphometric trait and environmental predictor using all sampled specimens. Statistical significance was assessed using Pearson's correlation test, with P-values adjusted for multiple comparisons using the Benjamini–Hochberg false discovery rate (FDR) correction.

| Environmental variable | Morphometric trait | Pearson r | P-value (adjusted) |
|---|---|---:|---:|
| Cation exchange capacity | Leaf length | 0.063 | 0.544 |
| Cation exchange capacity | Leaf width | 0.103 | 0.290 |
| Cation exchange capacity | Petiole length | 0.005 | 0.974 |
| Cation exchange capacity | Leaf base right angle | -0.010 | 0.961 |
| Cation exchange capacity | Leaf base left angle | -0.112 | 0.403 |
| Cation exchange capacity | Leaf apex angle | -0.121 | 0.209 |
| Cation exchange capacity | Calyx length | -0.183 | 0.087 |
| Cation exchange capacity | Calyx width | 0.127 | 0.273 |
| Cation exchange capacity | Corolla tube length | 0.722 | 0.000 |
| Cation exchange capacity | Pedicel length | 0.299 | 0.002 |
| Cation exchange capacity | Calyx teeth length | -0.161 | 0.171 |
| Cation exchange capacity | Calyx teeth width | 0.005 | 0.974 |
| Cation exchange capacity | Calyx teeth angle | 0.281 | 0.014 |
| Clay content | Leaf length | 0.119 | 0.209 |
| Clay content | Leaf width | 0.103 | 0.291 |
| Clay content | Petiole length | 0.215 | 0.010 |
| Clay content | Leaf base right angle | -0.163 | 0.183 |
| Clay content | Leaf base left angle | -0.196 | 0.090 |
| Clay content | Leaf apex angle | -0.116 | 0.235 |
| Clay content | Calyx length | 0.104 | 0.403 |
| Clay content | Calyx width | 0.022 | 0.870 |
| Clay content | Corolla tube length | 0.315 | 0.002 |
| Clay content | Pedicel length | 0.112 | 0.353 |
| Clay content | Calyx teeth length | -0.034 | 0.816 |
| Clay content | Calyx teeth width | 0.049 | 0.741 |
| Clay content | Calyx teeth angle | 0.079 | 0.596 |
| Coarse fragments (%) | Leaf length | 0.049 | 0.650 |
| Coarse fragments (%) | Leaf width | 0.030 | 0.795 |
| Coarse fragments (%) | Petiole length | 0.066 | 0.535 |
| Coarse fragments (%) | Leaf base right angle | -0.024 | 0.870 |
| Coarse fragments (%) | Leaf base left angle | -0.010 | 0.961 |
| Coarse fragments (%) | Leaf apex angle | 0.042 | 0.729 |
| Coarse fragments (%) | Calyx length | 0.194 | 0.062 |
| Coarse fragments (%) | Calyx width | 0.009 | 0.961 |
| Coarse fragments (%) | Corolla tube length | -0.229 | 0.033 |
| Coarse fragments (%) | Pedicel length | -0.079 | 0.535 |
| Coarse fragments (%) | Calyx teeth length | 0.148 | 0.209 |
| Coarse fragments (%) | Calyx teeth width | -0.001 | 0.994 |
| Coarse fragments (%) | Calyx teeth angle | -0.101 | 0.480 |
| Isothermality | Leaf length | 0.256 | 0.001 |

| Environmental variable | Morphometric trait | Pearson r | P-value (adjusted) |
|---|---|---|---|
| Isothermality | Leaf width | 0.105 | 0.253 |
| Isothermality | Petiole length | 0.116 | 0.209 |
| Isothermality | Leaf base right angle | -0.179 | 0.097 |
| Isothermality | Leaf base left angle | -0.136 | 0.240 |
| Isothermality | Leaf apex angle | -0.195 | 0.016 |
| Isothermality | Calyx length | -0.131 | 0.223 |
| Isothermality | Calyx width | 0.169 | 0.097 |
| Isothermality | Corolla tube length | 0.413 | 0.000 |
| Isothermality | Pedicel length | 0.324 | 0.000 |
| Isothermality | Calyx teeth length | -0.121 | 0.308 |
| Isothermality | Calyx teeth width | 0.088 | 0.480 |
| Isothermality | Calyx teeth angle | 0.306 | 0.006 |
| Mean diurnal range | Leaf length | 0.177 | 0.028 |
| Mean diurnal range | Leaf width | 0.076 | 0.447 |
| Mean diurnal range | Petiole length | 0.174 | 0.033 |
| Mean diurnal range | Leaf base right angle | -0.178 | 0.099 |
| Mean diurnal range | Leaf base left angle | -0.048 | 0.741 |
| Mean diurnal range | Leaf apex angle | -0.004 | 0.974 |
| Mean diurnal range | Calyx length | 0.256 | 0.006 |
| Mean diurnal range | Calyx width | 0.008 | 0.962 |
| Mean diurnal range | Corolla tube length | -0.451 | 0.000 |
| Mean diurnal range | Pedicel length | -0.136 | 0.209 |
| Mean diurnal range | Calyx teeth length | 0.185 | 0.086 |
| Mean diurnal range | Calyx teeth width | 0.035 | 0.802 |
| Mean diurnal range | Calyx teeth angle | -0.109 | 0.447 |
| Mean temp. driest quarter | Leaf length | -0.169 | 0.037 |
| Mean temp. driest quarter | Leaf width | -0.089 | 0.353 |
| Mean temp. driest quarter | Petiole length | -0.235 | 0.002 |
| Mean temp. driest quarter | Leaf base right angle | 0.167 | 0.135 |
| Mean temp. driest quarter | Leaf base left angle | 0.028 | 0.857 |
| Mean temp. driest quarter | Leaf apex angle | 0.057 | 0.582 |
| Mean temp. driest quarter | Calyx length | -0.329 | 0.000 |
| Mean temp. driest quarter | Calyx width | -0.074 | 0.535 |
| Mean temp. driest quarter | Corolla tube length | 0.346 | 0.000 |
| Mean temp. driest quarter | Pedicel length | 0.017 | 0.910 |
| Mean temp. driest quarter | Calyx teeth length | -0.339 | 0.000 |
| Mean temp. driest quarter | Calyx teeth width | -0.101 | 0.422 |
| Mean temp. driest quarter | Calyx teeth angle | 0.202 | 0.097 |
| Mean temp. wettest quarter | Leaf length | -0.194 | 0.014 |
| Mean temp. wettest quarter | Leaf width | -0.081 | 0.416 |
| Mean temp. wettest quarter | Petiole length | -0.229 | 0.003 |
| Mean temp. wettest quarter | Leaf base right angle | 0.184 | 0.091 |
| Mean temp. wettest quarter | Leaf base left angle | 0.052 | 0.722 |
| Mean temp. wettest quarter | Leaf apex angle | 0.115 | 0.209 |
| Mean temp. wettest quarter | Calyx length | -0.268 | 0.003 |
| Mean temp. wettest quarter | Calyx width | -0.106 | 0.353 |
| Mean temp. wettest quarter | Corolla tube length | 0.183 | 0.096 |

| Environmental variable | Morphometric trait | Pearson r | P-value (adjusted) |
|---|---|---|---|
| Mean temp. wettest quarter | Pedicel length | -0.086 | 0.465 |
| Mean temp. wettest quarter | Calyx teeth length | -0.287 | 0.003 |
| Mean temp. wettest quarter | Calyx teeth width | -0.130 | 0.264 |
| Mean temp. wettest quarter | Calyx teeth angle | 0.119 | 0.403 |
| Organic carbon stock | Leaf length | -0.070 | 0.494 |
| Organic carbon stock | Leaf width | -0.036 | 0.749 |
| Organic carbon stock | Petiole length | -0.009 | 0.958 |
| Organic carbon stock | Leaf base right angle | 0.038 | 0.802 |
| Organic carbon stock | Leaf base left angle | -0.044 | 0.759 |
| Organic carbon stock | Leaf apex angle | -0.042 | 0.729 |
| Organic carbon stock | Calyx length | 0.004 | 0.974 |
| Organic carbon stock | Calyx width | 0.029 | 0.849 |
| Organic carbon stock | Corolla tube length | -0.025 | 0.870 |
| Organic carbon stock | Pedicel length | 0.088 | 0.477 |
| Organic carbon stock | Calyx teeth length | 0.002 | 0.983 |
| Organic carbon stock | Calyx teeth width | -0.023 | 0.870 |
| Organic carbon stock | Calyx teeth angle | -0.075 | 0.615 |
| Precip. coldest quarter | Leaf length | -0.072 | 0.465 |
| Precip. coldest quarter | Leaf width | 0.021 | 0.861 |
| Precip. coldest quarter | Petiole length | -0.091 | 0.353 |
| Precip. coldest quarter | Leaf base right angle | 0.051 | 0.729 |
| Precip. coldest quarter | Leaf base left angle | -0.029 | 0.851 |
| Precip. coldest quarter | Leaf apex angle | -0.103 | 0.281 |
| Precip. coldest quarter | Calyx length | -0.327 | 0.000 |
| Precip. coldest quarter | Calyx width | 0.069 | 0.573 |
| Precip. coldest quarter | Corolla tube length | 0.648 | 0.000 |
| Precip. coldest quarter | Pedicel length | 0.315 | 0.000 |
| Precip. coldest quarter | Calyx teeth length | -0.216 | 0.033 |
| Precip. coldest quarter | Calyx teeth width | -0.043 | 0.751 |
| Precip. coldest quarter | Calyx teeth angle | 0.269 | 0.017 |
| Precip. seasonality | Leaf length | 0.116 | 0.209 |
| Precip. seasonality | Leaf width | 0.017 | 0.870 |
| Precip. seasonality | Petiole length | 0.176 | 0.032 |
| Precip. seasonality | Leaf base right angle | -0.121 | 0.319 |
| Precip. seasonality | Leaf base left angle | 0.042 | 0.754 |
| Precip. seasonality | Leaf apex angle | 0.081 | 0.423 |
| Precip. seasonality | Calyx length | 0.369 | 0.000 |
| Precip. seasonality | Calyx width | -0.014 | 0.921 |
| Precip. seasonality | Corolla tube length | -0.625 | 0.000 |
| Precip. seasonality | Pedicel length | -0.243 | 0.010 |
| Precip. seasonality | Calyx teeth length | 0.251 | 0.010 |
| Precip. seasonality | Calyx teeth width | 0.097 | 0.447 |
| Precip. seasonality | Calyx teeth angle | -0.269 | 0.017 |
| Precip. warmest quarter | Leaf length | 0.141 | 0.097 |
| Precip. warmest quarter | Leaf width | 0.118 | 0.201 |
| Precip. warmest quarter | Petiole length | 0.276 | 0.000 |
| Precip. warmest quarter | Leaf base right angle | -0.204 | 0.053 |

| Environmental variable | Morphometric trait | Pearson r | P-value (adjusted) |
|---|---|---|---|
| Precip. warmest quarter | Leaf base left angle | 0.045 | 0.749 |
| Precip. warmest quarter | Leaf apex angle | 0.041 | 0.729 |
| Precip. warmest quarter | Calyx length | 0.382 | 0.000 |
| Precip. warmest quarter | Calyx width | 0.096 | 0.417 |
| Precip. warmest quarter | Corolla tube length | -0.314 | 0.001 |
| Precip. warmest quarter | Pedicel length | -0.081 | 0.491 |
| Precip. warmest quarter | Calyx teeth length | 0.325 | 0.000 |
| Precip. warmest quarter | Calyx teeth width | 0.047 | 0.741 |
| Precip. warmest quarter | Calyx teeth angle | -0.201 | 0.097 |
| Silt content | Leaf length | 0.074 | 0.470 |
| Silt content | Leaf width | 0.096 | 0.337 |
| Silt content | Petiole length | -0.026 | 0.834 |
| Silt content | Leaf base right angle | 0.047 | 0.749 |
| Silt content | Leaf base left angle | -0.077 | 0.573 |
| Silt content | Leaf apex angle | -0.123 | 0.209 |
| Silt content | Calyx length | -0.085 | 0.486 |
| Silt content | Calyx width | 0.197 | 0.058 |
| Silt content | Corolla tube length | 0.586 | 0.000 |
| Silt content | Pedicel length | 0.264 | 0.006 |
| Silt content | Calyx teeth length | -0.150 | 0.209 |
| Silt content | Calyx teeth width | 0.098 | 0.448 |
| Silt content | Calyx teeth angle | 0.300 | 0.008 |
| Soil bulk density | Leaf length | 0.079 | 0.447 |
| Soil bulk density | Leaf width | 0.050 | 0.644 |
| Soil bulk density | Petiole length | -0.057 | 0.597 |
| Soil bulk density | Leaf base right angle | -0.016 | 0.921 |
| Soil bulk density | Leaf base left angle | 0.079 | 0.563 |
| Soil bulk density | Leaf apex angle | 0.019 | 0.870 |
| Soil bulk density | Calyx length | -0.066 | 0.599 |
| Soil bulk density | Calyx width | -0.060 | 0.644 |
| Soil bulk density | Corolla tube length | -0.027 | 0.861 |
| Soil bulk density | Pedicel length | -0.154 | 0.171 |
| Soil bulk density | Calyx teeth length | -0.097 | 0.448 |
| Soil bulk density | Calyx teeth width | -0.098 | 0.448 |
| Soil bulk density | Calyx teeth angle | 0.091 | 0.535 |
| Soil nitrogen | Leaf length | -0.063 | 0.544 |
| Soil nitrogen | Leaf width | 0.003 | 0.981 |
| Soil nitrogen | Petiole length | 0.059 | 0.583 |
| Soil nitrogen | Leaf base right angle | -0.066 | 0.639 |
| Soil nitrogen | Leaf base left angle | -0.072 | 0.598 |
| Soil nitrogen | Leaf apex angle | -0.033 | 0.781 |
| Soil nitrogen | Calyx length | -0.006 | 0.974 |
| Soil nitrogen | Calyx width | 0.104 | 0.403 |
| Soil nitrogen | Corolla tube length | 0.430 | 0.000 |
| Soil nitrogen | Pedicel length | 0.251 | 0.010 |
| Soil nitrogen | Calyx teeth length | -0.047 | 0.746 |
| Soil nitrogen | Calyx teeth width | 0.017 | 0.919 |

| Environmental variable | Morphometric trait | Pearson r | P-value (adjusted) |
| --- | --- | --- | --- |
| Soil nitrogen | Calyx teeth angle | 0.051 | 0.749 |

**TABLE S5. Membership probabilities of each individual to its assigned morphotype according to DAPC analysis**. Ind. = Individual. Localities described for specimens of considerably mixed memberships probabilities.

| Ind. | bahia | capitata | hydrangeiformis | Voucher | Locality |
|---|---|---|---|---|---|
| 1 | 0.999 | 0.000 | 0.000 | BHCB166937GiacominLL1984 | |
| 2 | 0.999 | 0.000 | 0.000 | BHCB104358AmorimAM5263 | |
| 3 | 0.999 | 0.000 | 0.000 | BHCB112709JardimJG4373 | |
| 4 | 0.999 | 0.000 | 0.000 | RB554059JardimAB83 | |
| 5 | 0.999 | 0.000 | 0.000 | RB317933SantosEB171 | |
| 6 | 0.999 | 0.000 | 0.000 | MO2659829FiaschiP1126 | |
| 7 | 0.999 | 0.000 | 0.000 | SP370911AmorimAM3586 | |
| 8 | 0.999 | 0.000 | 0.000 | NY684452ThomasWW12025 | |
| 9 | 0.999 | 0.000 | 0.000 | NY684453ThomasWW11383 | |
| 10 | 0.999 | 0.000 | 0.000 | NY684489MattosSilvaLA3603 | |
| 11 | 0.999 | 0.000 | 0.000 | NY684557ThomasWW11207 | |
| 12 | 0.999 | 0.000 | 0.000 | NY00864748AmorimAM4353 | |
| 13 | 0.999 | 0.000 | 0.000 | NY1280984CarvalhoAMV4445 | |
| 14 | 0.999 | 0.000 | 0.000 | NY01805847JardimJG4810 | |
| 15 | 0.999 | 0.000 | 0.000 | NY01805994ReginatoM217 | |
| 16 | 0.999 | 0.000 | 0.000 | NY01806248AmorimAM6320 | |
| 17 | 0.999 | 0.000 | 0.000 | NY01806263AmorimAM4458 | |
| 18 | 0.999 | 0.000 | 0.000 | CEN51879OliveiraJS23 | |
| 19 | 0.999 | 0.000 | 0.000 | CEPEC116154BorgesRAX624 | |
| 20 | 0.999 | 0.000 | 0.000 | CEPEC128639LiuthHS67 | |
| 21 | 0.999 | 0.000 | 0.000 | CEPEC123912DaneuL43 | |
| 22 | 0.999 | 0.000 | 0.000 | CEPEC126284PerdizRO410 | |
| 23 | 0.999 | 0.000 | 0.000 | CEPEC126268PerdizRO394 | |
| 24 | 0.935 | 0.000 | 0.064 | CEPEC110870AmorimAM5581 | |
| 25 | 0.999 | 0.000 | 0.000 | CEPEC110469AmorimAM5425 | |

| Ind. | bahia | capitata | hydrangeiformis | Voucher | Locality |
|---|---|---|---|---|---|
| 26 | 0.999 | 0.000 | 0.000 | CEPEC128935PerdizRO797 | |
| 27 | 0.999 | 0.000 | 0.000 | CEPEC99982MattosSilvaLA4816 | |
| 28 | 0.999 | 0.000 | 0.000 | CEPEC126516PerdizRO483 | |
| 29 | 0.999 | 0.000 | 0.000 | CEPEC109966JardimJG4844 | |
| 30 | 0.999 | 0.000 | 0.000 | CEPEC106230ThomasWW14118 | |
| 31 | 0.000 | 0.965 | 0.034 | MBML003989HoffmannWA233 | |
| 32 | 0.000 | 0.942 | 0.057 | MBML004037VimercatJM73 | |
| 33 | 0.000 | 0.920 | 0.079 | MBML001723BoudetFernandesHQ1819 | |
| 34 | 0.000 | 0.993 | 0.006 | MBML037629ForzzaRC5390 | |
| 35 | 0.000 | 0.973 | 0.026 | MBML006595BausenE58 | |
| 36 | 0.000 | 0.993 | 0.006 | MBML001725BoudetFernandesHQ1811 | |
| 37 | 0.000 | 0.690 | 0.309 | MBML018950VervloetRR1591 | |
| 38 | 0.000 | 0.997 | 0.002 | MBML025334KollmannL8532 | |
| 39 | 0.000 | 0.975 | 0.024 | MBML012104BausenE140 | |
| 40 | 0.000 | 0.998 | 0.001 | MBML001717BoudetFernandesHQ1790 | |
| 41 | 0.000 | 0.848 | 0.150 | MBML004046VimercatJM170 | Espírito Santo, Santa Teresa, Rio Saltinho. |
| 42 | 0.000 | 0.943 | 0.056 | MBML008222KollmannL1368 | |
| 43 | 0.000 | 0.983 | 0.016 | MBML023929KollmannL7282 | |
| 44 | 0.000 | 0.995 | 0.004 | MBML028186BrittoRC63 | |
| 45 | 0.000 | 0.993 | 0.006 | MBML037990AssisAM1337 | |
| 46 | 0.000 | 0.979 | 0.020 | MBML039849AssisAM1139 | |
| 47 | 0.000 | 0.694 | 0.305 | MBML481216FontanaAP2777 | Espírito Santo, Santa Teresa, APA Goiapaba-açu. |
| 48 | 0.000 | 0.944 | 0.055 | MBML042116FiaschiP3527 | |
| 49 | 0.000 | 0.976 | 0.023 | MBML011947ChamasC121 | |
| 50 | 0.000 | 0.995 | 0.004 | BHCB88681MotaRC2246 | |
| 51 | 0.000 | 0.995 | 0.004 | BHCB163875GiorniVTsn | |
| 52 | 0.000 | 0.999 | 0.000 | BHCB171745GianerriniAC16 | |

| Ind. | bahia | capitata | hydrangeiformis | Voucher | Locality |
|---|---|---|---|---|---|
| 53 | 0.000 | 0.998 | 0.001 | BHCB208726MessederJVS82 | |
| 54 | 0.000 | 0.743 | 0.256 | BHCB63824MotaRC542 | Minas Gerais, Mariana, Mina da Samitri. |
| 55 | 0.000 | 0.987 | 0.012 | BHCB146029RezendeSG3863 | |
| 56 | 0.000 | 0.172 | 0.827 | BHCB105183MotaRC3161 | Minas Gerais, Mariana, Serra do Caraça. |
| 57 | 0.000 | 0.999 | 0.000 | BHCB60086BrinaAEsn | |
| 58 | 0.000 | 0.999 | 0.000 | BHCB53921TeixeiraWAsn | |
| 59 | 0.000 | 0.990 | 0.009 | BHCB186577CastroR2601 | |
| 60 | 0.000 | 0.994 | 0.005 | BHCB148507GiacominLL898 | |
| 61 | 0.000 | 0.843 | 0.156 | BHCB144959GiacominLL847 | São Paulo, Bananal, Estação Ecológica de Bananal. |
| 62 | 0.000 | 0.999 | 0.000 | BHCB209638NadruzM3786 | |
| 63 | 0.000 | 0.081 | 0.918 | BHCB57657SalinoA6239 | |
| 64 | 0.000 | 0.170 | 0.829 | BHCB174046StehmannJR6355 | Minas Gerais, Santa Barbara, Conceiçao do Rio Acima. |
| 65 | 0.000 | 0.937 | 0.062 | BHCB152244GiacominLL1209 | Espírito Santo, Santa Teresa, Estação Biológica de Santa Lúcia. |
| 66 | 0.000 | 0.998 | 0.001 | BHCB157209OliveiraVE191 | |
| 67 | 0.000 | 0.988 | 0.011 | BHCB200593MartinelliG19860 | |
| 68 | 0.000 | 0.961 | 0.038 | BHCB77040SilvaLsn | |
| 69 | 0.000 | 0.983 | 0.016 | RB81821MarkgrafF10495 | |
| 70 | 0.000 | 0.999 | 0.000 | RB526119BorgesR1203 | |
| 71 | 0.000 | 0.999 | 0.000 | RB847062FernandezE1092 | |
| 72 | 0.000 | 0.987 | 0.012 | RB866339CarvalhoRGG24 | |
| 73 | 0.000 | 0.999 | 0.000 | RB821752NadruzM3773 | |
| 74 | 0.000 | 0.999 | 0.000 | RB460017NadruzM2105 | |
| 75 | 0.000 | 0.998 | 0.001 | RB801486KulkampJ1103 | |
| 76 | 0.000 | 0.858 | 0.141 | RB800925BaezC1854 | Rio de Janeiro, Itatiaia, Parque Nacional do Itatiaia. |
| 77 | 0.000 | 0.875 | 0.124 | RB729903MartinelliG19860 | Rio de Janeiro, Rio Claro, Parque Estadual do Cunhambebe. |
| 78 | 0.000 | 0.999 | 0.000 | RB138762SucreD2554 | |
| 79 | 0.000 | 0.990 | 0.009 | RB654535CardosoLJT1473 | |

| Ind. | bahia | capitata | hydrangeiformis | Voucher | Locality |
|---|---|---|---|---|---|
| 80 | 0.000 | 0.999 | 0.000 | RB488137BorgesR962 | |
| 81 | 0.000 | 0.851 | 0.148 | RB777600NadruzM3460 | Minas Gerais, Fervedouro, Parque Estadual Serra do Brigadeiro. |
| 82 | 0.000 | 0.996 | 0.003 | RB680998FerreiraCDM424 | |
| 83 | 0.000 | 0.971 | 0.028 | RB645658LirioEJ1079 | |
| 84 | 0.000 | 0.997 | 0.002 | RB643273CardosoLJT1186 | |
| 85 | 0.000 | 0.973 | 0.026 | RB268020HatschbachG49870 | |
| 86 | 0.000 | 0.996 | 0.003 | RB680306LeoniLS149 | |
| 87 | 0.000 | 0.993 | 0.006 | RB680303LeoniLS2352 | |
| 88 | 0.000 | 0.993 | 0.006 | RB870945CarvalhoRGG34 | |
| 89 | 0.000 | 0.999 | 0.000 | RB431165PifanoDS189 | |
| 90 | 0.000 | 0.976 | 0.023 | RB736441LeoniLS5231 | |
| 91 | 0.000 | 0.999 | 0.000 | RB48082HeringerEP460 | |
| 92 | 0.000 | 0.999 | 0.000 | RB736444LeoniLS6673 | |
| 93 | 0.000 | 0.941 | 0.058 | RB553367IvanauskasNM6360 | |
| 94 | 0.000 | 0.895 | 0.104 | RB186775MartinelliG1115 | São Paulo, Bananal, Parque Nacional da Serra da Bocaina. |
| 95 | 0.000 | 0.993 | 0.006 | RB207446MartinelliG7777 | |
| 96 | 0.000 | 0.900 | 0.099 | RB438923LimaHC5517 | |
| 97 | 0.000 | 0.999 | 0.000 | RB460725NadruzM2181 | |
| 98 | 0.000 | 0.682 | 0.317 | RB628148FiaschiP3773 | Rio de Janeiro, Casimiro de Abreu, Reserva Biológica União. |
| 99 | 0.000 | 0.987 | 0.012 | RB870945CarvalhoRGG34 | |
| 100 | 0.000 | 0.999 | 0.000 | RB475622NadruzM2248 | |
| 101 | 0.000 | 0.995 | 0.004 | RB25993PortoC1879 | |
| 102 | 0.000 | 0.921 | 0.078 | RB88993DuarteAP4109 | |
| 103 | 0.000 | 0.997 | 0.002 | SP310763AlbuquerqueLB3 | |
| 104 | 0.000 | 0.979 | 0.020 | SP303135RossiL1584 | |
| 105 | 0.000 | 0.998 | 0.001 | SP192431CustodioFilhoA1284 | |
| 106 | 0.000 | 0.999 | 0.000 | SP79377KuhlmannM1754 | |

| Ind. | bahia | capitata | hydrangeiformis | Voucher | Locality |
|---|---|---|---|---|---|
| 107 | 0.000 | 0.995 | 0.004 | SP197600CustodioFilhoA2005 | |
| 108 | 0.000 | 0.971 | 0.028 | SP56411HandroO744 | |
| 109 | 0.000 | 0.994 | 0.005 | SP48577LimaASsn | |
| 110 | 0.049 | 0.944 | 0.005 | SP15183EdwallG15183 | |
| 111 | 0.000 | 0.999 | 0.000 | MBM227974TeixeiraEMsn | |
| 112 | 0.000 | 0.999 | 0.000 | US2613938CarautaJPP716 | |
| 113 | 0.000 | 0.999 | 0.000 | US1906065WilliamsLO7928 | |
| 114 | 0.000 | 0.902 | 0.097 | MBM440676VoltzRR2752 | |
| 115 | 0.000 | 0.072 | 0.927 | MBML037156ForzzaRC5429 | Espírito Santo, Castelo, Parque Estadual do Forno Grande. |
| 116 | 0.000 | 0.079 | 0.920 | MBML013774DemunerV1480 | Espírito Santo, Santa Teresa, Alto São Lourenço. |
| 117 | 0.000 | 0.512 | 0.487 | MBML022321KollmannL6668 | Espírito Santo, Castelo, Parque Estadual do Forno Grande. |
| 118 | 0.000 | 0.058 | 0.941 | MBML037356FragaCN2223 | |
| 119 | 0.000 | 0.100 | 0.899 | BHCB162191GouveaYF08 | Minas Gerais, Simonésia, RPPN Mata do Sossego. |
| 120 | 0.000 | 0.012 | 0.987 | BHCB175530FalcaoBF2 | |
| 121 | 0.000 | 0.012 | 0.987 | BHCB102229VianaPL2498 | |
| 122 | 0.987 | 0.000 | 0.012 | BHCB166905GiacominLL1952 | |
| 123 | 0.000 | 0.129 | 0.870 | BHCB3288GrandiTSM756 | Minas Gerais, Juiz de Fora. |
| 124 | 0.820 | 0.000 | 0.179 | BHCB91115StehmannJR4024 | Minas Gerais, Santa Maria do Salto. |
| 125 | 0.000 | 0.853 | 0.146 | RB561684SalgadoCS228 | Rio de Janeiro, Miguel Pereira. |
| 126 | 0.000 | 0.021 | 0.978 | RB666202BaezC1097 | |
| 127 | 0.000 | 0.003 | 0.996 | RB762413VerdiM7505 | |
| 128 | 0.000 | 0.000 | 0.999 | RB296553AraujoIA84 | |
| 129 | 0.000 | 0.004 | 0.995 | RB125484DuarteAPs.n. | |
| 130 | 0.000 | 0.010 | 0.989 | RB251984MartinelliG11964 | |
| 131 | 0.000 | 0.006 | 0.993 | RB250418MartinelliG11725 | |
| 132 | 0.000 | 0.000 | 0.999 | RB406987SilvaBR1338 | |
| 133 | 0.000 | 0.000 | 0.999 | RB27090LimaS200 | |

| Ind. | bahia | capitata | hydrangeiformis | Voucher | Locality |
|---|---|---|---|---|---|
| 134 | 0.000 | 0.353 | 0.646 | RB542103BragaJMA3645 | |
| 135 | 0.000 | 0.273 | 0.726 | RB22603DuckeAsn | Rio de Janeiro, Tijuca, Bico do Papagaio. |
| 136 | 0.000 | 0.953 | 0.046 | RB8337PortoC658 | |
| 137 | 0.000 | 0.000 | 0.999 | RB665784FerreiraCDM370 | |
| 138 | 0.000 | 0.827 | 0.172 | RB438987NadruzM563 | Rio de Janeiro, Nova Friburgo, Macaé de Cima. |
| 139 | 0.000 | 0.499 | 0.500 | SPF184414KriegerPL10763 | Minas Gerais, Juiz de Fora. |
| 140 | 0.000 | 0.940 | 0.059 | HUEM6285SakuraguiCM921 | |
| 141 | 0.000 | 0.292 | 0.707 | MBM342700KriegerL11806 | Minas Gerais, Juiz de Fora. |
| 142 | 0.000 | 0.064 | 0.935 | MBM342701KriegerL9219 | |
| 143 | 0.000 | 0.855 | 0.144 | VIES21494ZorzanelliJPF110 | Espírito Santo, Iúna, Serra do Valentim. |
| 144 | 0.000 | 0.104 | 0.895 | MBM169647HatschbachG1994 | Espírito Santo, Domingos Martins, rio Jucu Braço Sul. |
| 145 | 0.000 | 0.802 | 0.197 | MBM255795HatschbachG71906 | Espírito Santo, Castelo, Caxixe Quente. |
| 146 | 0.000 | 0.003 | 0.996 | CEPEC63618HatschbachC61193 | |

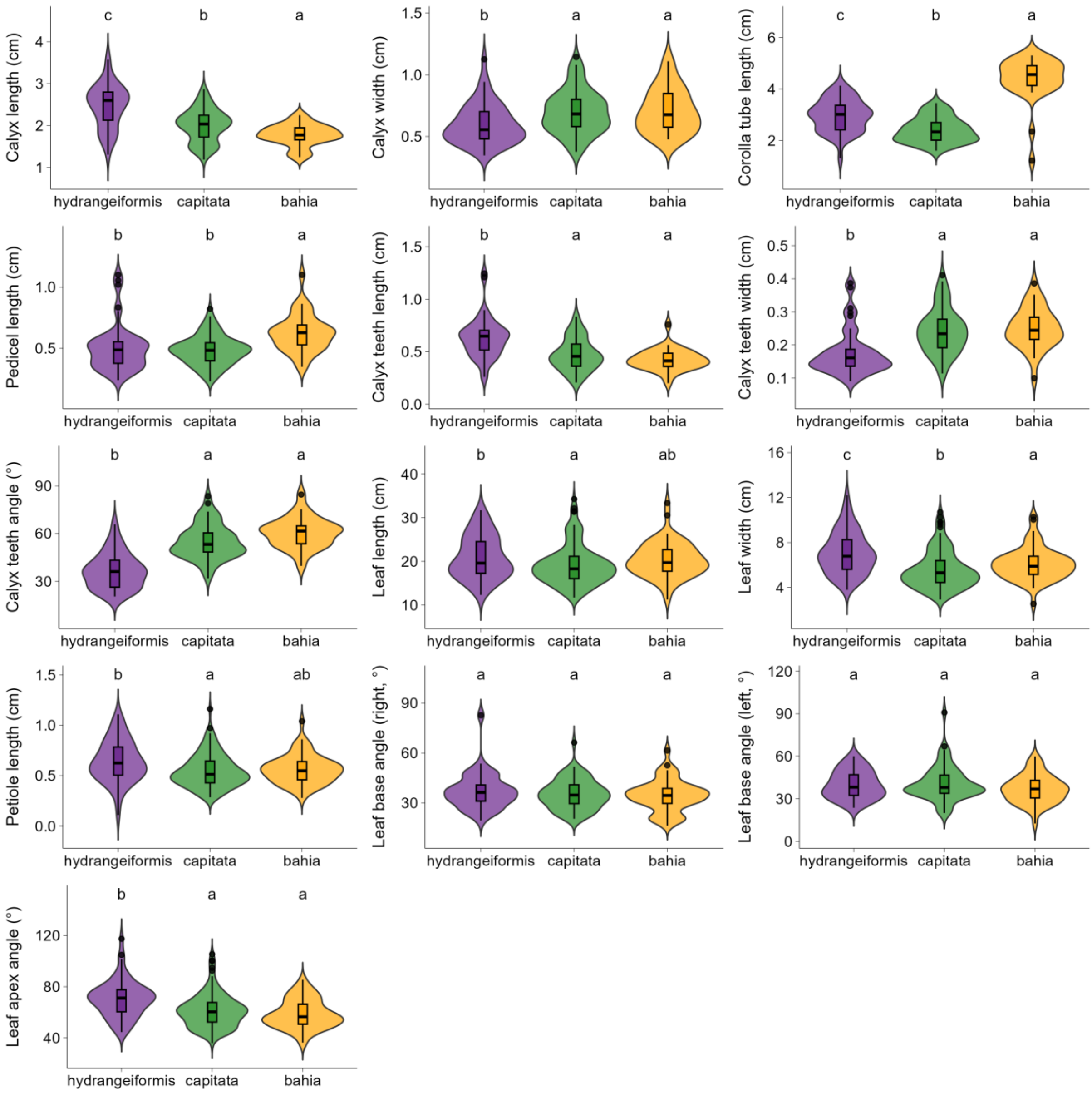


**FIGURE S1. Violin plots for all morphometric characters analyzed.** Graphs illustrate the distribution and variability of each morphological trait across morphotypes, highlighting differences in central tendency and dispersion. Different letters (a,b,c) denote significant differences among morphotypes based on ANOVA or Kruskal–Wallis tests followed by post hoc comparisons ($P < 0.05$).
